# Bacteriophage infection drives loss of β-lactam resistance in methicillin-resistant *Staphylococcus aureus*

**DOI:** 10.1101/2024.08.19.608629

**Authors:** My Tran, Angel J. Hernandez Viera, Patricia Q. Tran, Erick Nilsen, Lily Tran, Charlie Y. Mo

## Abstract

Bacteriophage (phage) therapy is a promising means to combat drug-resistant bacterial pathogens. Infection by phage can select for mutations in bacterial populations that confer resistance against phage infection. However, resistance against phage can yield evolutionary trade-offs of biomedical relevance. Here we report the discovery that infection by certain staphylococcal phages sensitizes different strains of methicillin-resistant *Staphylococcus aureus* (MRSA) to β-lactams, a class of antibiotics against which MRSA is typically resistant. MRSA cells that survive infection by these phages display significant reductions in minimal inhibitory concentration against different β-lactams compared to uninfected bacteria. Transcriptomic profiling reveals that these evolved MRSA strains possess highly modulated transcriptional profiles, where numerous genes involved in *S. aureus* virulence were downregulated. Phage-treated MRSA exhibited attenuated virulence phenotypes in the form of reduced hemolysis and clumping. Despite sharing similar phenotypes, whole-sequencing analysis revealed that the different MRSA strains evolved unique genetic profiles during infection. These results suggest complex evolutionary trajectories in MRSA during phage predation and open up new possibilities to reduce drug resistance and virulence in MRSA infections.

---

*Staphylococcus aureus* is one of the most notorious and widespread bacterial pathogens, responsible for hundreds of thousands of severe infections worldwide every year. Methicillin-resistant *S. aureus* (MRSA) poses a particular clinical threat, as MRSA infections increase mortality, morbidity, and hospital stay, as compared to those caused by methicillin-sensitive *S. aureus* (MSSA).^1^ Part of MRSA’s notoriety stems from its strong resistance against the β-lactam family of antibiotics, such as penicillins and cephalosporins, which inhibit the activity of transpeptidase enzymes during peptidoglycan synthesis in bacterial cell walls.^2^ β-lactams are one of the most commonly prescribed drug classes, with many designated as “Critically Important” antimicrobials by the World Health Organization.^3^ Thus, MRSA infections pose a considerable public health risk as they are notoriously difficult to treat and are widespread in communities and hospital settings. Indeed, in 2019, MRSA alone accounted for more than 100,000 deaths attributable to drug-resistant infections worldwide.^4^

A chief mediator of β-lactams resistance in MRSA is the SCCmec cassette, a mobile genetic element that carries the resistance gene *mecA.* MecA encodes for the penicillin-binding protein 2A (PBP2a), a transpeptidase that has a low affinity for β-lactams.^5^ This lower affinity permits PBP2a to participate in peptidoglycan synthesis even in the presence of β-lactams, ultimately resulting in cell survival. In addition to *mecA*, numerous MRSA strains also carry β-lactamases, such as BlaZ, that degrade β-lactams, thus further contributing to drug resistance. Together, these mechanisms can severely limit treatment options against MRSA, with current clinical treatment options relying primarily on vancomycin and daptomycin.^6^ Both vancomycin and daptomycin are last resort antibiotics against MRSA, and a major concern is the increasing resistance of MRSA to these drugs.^7,8^ Developing solutions to combat MRSA is a major focus in academia and industry.

Due to its drug resistance and clinical burden, *S. aureus* is a prime candidate for alternative antimicrobial treatments, suchas bacteriophage (phage) therapy. Phages are viruses that infect and kill bacteria, posing one of the greatest existential threats to bacterial communities, with some estimates suggesting that 40% of all bacterial mortality worldwide is caused by phage predation.^9^ In phage therapy, lytic phages are administered to kill the bacterial pathogen(s) causing an infection. Phages offer certain advantages over traditional antibiotics: they are highly specific to their hosts by reducing off-target killing; they self-amplify and evolve, enabling the rapid generation of new phage variants with improved activities; and they are generally regarded as safe, as toxicity has been reported only in extremely rare cases in animals and patients.^10^ Indeed, against *S. aureus* infections, over a dozen promising case studies and clinical trials have been reported in the past decade.^11^

Despite these advances, routine use of phage therapy is still met with challenges. Chief among these is the inevitable rise of phage resistance, as phage predation exerts a strong selective pressure on bacterial populations. According to one meta-analysis that focused on phage therapy outcomes, resistance against phage evolved in 75 percent of human clinical cases in which the evolution of resistance was monitored.^12^ Mutations represent a chief pathway by which bacteria evolve resistance against phage. To date, the best characterized phage resistance mutations involve alterations on cell surface receptor molecules that mediate phage attachment. In many bacteria, these receptors are often proteins or sugar moieties, which are recognized by phage proteins.^11,13,14^ For example, in *Escherichia coli*, mutations in the cell-wall protein LamB confer resistance against *lambda* phage infection, while in *S. aureus*, mutations that modify wall teichoic acid (WTA) have been shown to limit phage infection.^14,15^ Complicating the picture, studies have revealed that a plethora of additional host mechanisms, including dedicated anti-phage defense systems, can impact the evolution of resistance against phage.^16,17^ These problems highlight the importance of developing phage treatment strategies that minimize or capitalize on the evolution of phage resistance.^18^

A unique aspect of phage therapy is the possibility to exploit evolutionary trade-offs to combat resistant pathogens. A genetic trade-off is defined as an evolved trait that confers a fitness advantage against a particular selective pressure at the expense of reduced fitness against an unselected pressure. Across many different species of bacteria, such trade-offs have been shown to occur between phage resistance and antibiotic resistance (Figure 1A). Phages that bind to a virulence factor or mechanism for antibiotic resistance in the target bacteria are predicted to exert a strong selection pressure on the bacteria to mutate or downregulate the phage-binding target. These changes would confer protection against phage infection but could in turn reduce the resistance or virulence in the bacterium. As an example, in *P. aeruginosa,* infection by the phage OMKO1, which binds to the outer membrane protein M OprM of MexAB- and MexXY-OprM efflux pumps, drives the evolution of mutations in those genes, leading to the re-sensitization of phage-resistant *P. aeruginosa* mutants to antibiotics.^10,19^

**Figure 1.**
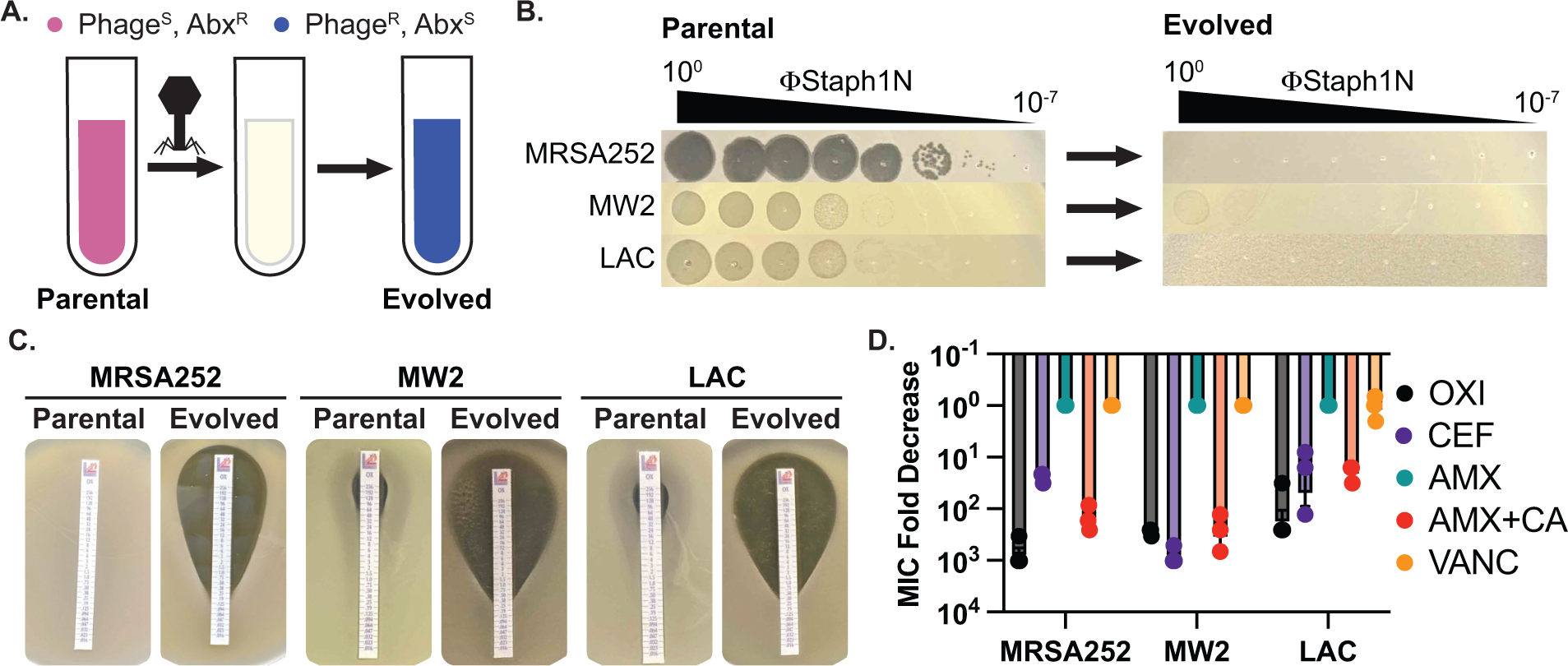
Infection by bacteriophage ΦStaph1N drives the loss of β-lactam resistance in MRSA. **A.** Schematic of the experimental setup. Drug-resistant (Abx^R^), phage-sensitive (Phage^S^) bacterial cultures are infected with phage. The population of infected cells is passaged and allowed to recover. The surviving cell population is resistant to phage infection (Phage^R^) but has evolved sensitivity to antibiotics (Abx^S^). **B.** ΦStaph1N infects MRSA strains MRSA252, MW2, and LAC (left panel). Following infection with ΦStaph1N, evolved cultures of the three MRSA strains are resistant to ΦStaph1N (right panel). **C.** ΦStaph1N-treated, evolved MRSA strains show significant loss of resistance against oxacillin, compared to the parental strains. Loss of resistance is indicated by the area of bacterial clearance surrounding the antibiotic resistance strip. **D.** ΦStaph1N treatment causes loss resistance against different β-lactams. Plotted are the fold reductions of minimal inhibitory concentration (MIC) between treated and mock-treated cells. OXI = oxacillin; CEF = cefazolin; AMX = amoxicillin; AMX+CA = amoxicillin & clavulanic acid; VANC = vancomycin.

Little is known about how phage resistance can mediate genetic trade-offs in MRSA. Previous work has shown that phage resistance in *S. aureus* can proceed through genetic mutations that are not directly involved in phage binding. For example, studies by Berryhill and colleagues demonstrated that phage infection of *S. aureus* Newman, an MSSA strain, can select for mutations in *femA*, which is a cytoplasmic enzyme that catalyzes the formation of the pentaglycine bridge of peptidoglycans in *S. aureus*.^20^ Rather than serving as a direct phage receptor molecule, *femA* maintains the integrity of the cell wall, which in turn could be vital for WTA maturation and phage attachment.^13,21^ Interestingly, a consequence of these *femA* mutations is increased sensitivity against antibiotics. We therefore asked how phage resistance might impact the physiology of drug-resistant MRSA, with the hopes of identifying genetic trade-offs of potential biomedical relevance.

In the work reported here, we show that infection by staphylococcal phages causes MRSA strains to evolve sensitivity to different types of β-lactam antibiotics and attenuate virulence phenotypes. We found that this loss of resistance and virulence is associated with distinct mutational profiles distinct in each MRSA strain, and that phage-treated, evolved MRSA populations display significant transcriptome remodeling. Unexpectedly, we also discovered a mutant phage with higher activity and a broader host range against MRSA. Findings from our work can help in the development of phage therapies that reduce drug resistance and virulence in pathogenic bacteria.

## Results

### Identification of ΦStaph1N with activity against multiple MRSA strains

For our studies, we focused on three MRSA strains – MRSA252 (USA200), MW2 (USA400), and LAC (USA300). All three MRSA strains are pathogenic isolates implicated in human disease and are used as representative examples for studying MRSA.^22–24^ To test for phage susceptibility, we performed plaquing assays with a panel of staphylococcal phages (Table S1, Figure S1). Of the phages tested, one phage, ΦStaph1N, which belongs to the *Kayvirus* genus, formed plaques on all three MRSA strains (Figure 1B).^21,25^ Yet, despite its ability to infect all three strains, ΦStaph1N infection was unable to eradicate MRSA cultures. Both MW2 and LAC displayed incomplete lysis in liquid culture at multiplicities of infection (MOIs) of 0.1 or lower (Figure S2). Furthermore, infected cultures of all three MRSA strains could recover back to high cell density after passaging one percent of the culture into fresh media following 24 hours of initial infection. These results suggest that infection by ΦStaph1N selects for resistant mutants that could sweep the population. Indeed, ΦStaph1N was unable to form plaques on recovered MRSA cultures that survived in the initial ΦStaph1N infection (Figure 1B).

### Resistance against ΦStaph1N infection sensitizes MRSA against β-lactams

Because both phages and β-lactams interface with the bacterial cell wall, we hypothesized that resistance against ΦStaph1N infection could cause a trade-off in β-lactam resistance in MRSA even in the presence of PBP2a and BlaZ. We first tested the β-lactam sensitivity of the parental MRSA252, MW2, and LAC strains. As expected, all three strains displayed high minimal inhibitory concentrations (MICs) of ≥48 µg/mL against the β-lactams oxacillin (OXA), cefazolin (CEF), amoxicillin (AMX), and amoxicillin & clavulanic acid (AMX+CA), visually indicated by their ability to form lawns surrounding antibiotic strips (Figure 1C, Table S2). The strains were sensitive to vancomycin (VANC; MICs = 1.5 µg/mL), which inhibits cell wall synthesis through a different mechanism than β-lactams. Strikingly, phage-resistant MRSA that survived ΦStaph1N infection displayed a strong reduction in resistance against OXA, CEF, and AMX+CA, with fold reductions in MIC between 10 and 1000-fold (Figure 1D); no change in MIC was observed with VANC or with AMX alone. These results show at a phenotypic level that ΦStaph1N-resistant MRSA loses resistance towards most β-lactams.

We next asked whether this loss of β-lactam resistance depended on the MOI of ΦStaph1N. We infected the three MRSA strains with ΦStaph1N at MOIs ranging from 10^−2^ to 10^−5^, isolated the surviving MRSA cells, and tested their MIC against oxacillin (Table S2). For MRSA252, we still observed a ∼3-order of magnitude fold reduction of MIC at an MOI of 10^−5^. With MW2, the reduction of MIC was markedly decreased with lower phage levels, showing no significant loss at MOIs of 10^−3^ or lower. For LAC, two replicates displayed a reduction of MIC by an order of magnitude at an MOI of 10^−4^, while the third replicate did not display any change. These results show that for MRSA252, ΦStaph1N MOIs as low as 10^−5^ can still drive the loss of resistance, while for MW2 and LAC, higher MOIs of phage are needed to ensure the same outcome of reduced β-lactam resistance.

### Discovery of a mutant ΦStaph1N with enhanced activity against MRSA

For ΦStaph1N, we noticed that while the phage could plaque on all three MRSA strains, its plaque-forming efficiency was reduced on the MW2 and LAC strains (Figure 1B, Figure S1). ΦStaph1N plaques on MW2 and LAC bacterial lawns were hazy, and the overall efficiency of plaquing was approximately two orders of magnitude less than that on MRSA252. Unexpectedly, we consistently observed smaller, clear plaques arising in the larger, hazy plaques of LAC (Figure S3A); notably this did not appear in MW2. We hypothesized that these clear plaques were caused by a mutant form of ΦStaph1N that evolved higher lytic activity. We isolated phage clones from these single plaques and tested their activity against MRSA. This mutant phage, which we called Evo2, plaques on LAC and MW2 strains with higher efficiency, displaying comparable plaquing to MRSA252 (Figure 2A). In growth experiments, we further observed that Evo2 lyses MRSA cultures at lower MOIs compared to ΦStaph1N (Figure S2). Evo2 exhibits lytic activity against MW2 and LAC even at an MOI of 10^−4^, a concentration at which ΦStaph1N does not show any detectable activity against the two strains.

**Figure 2.**
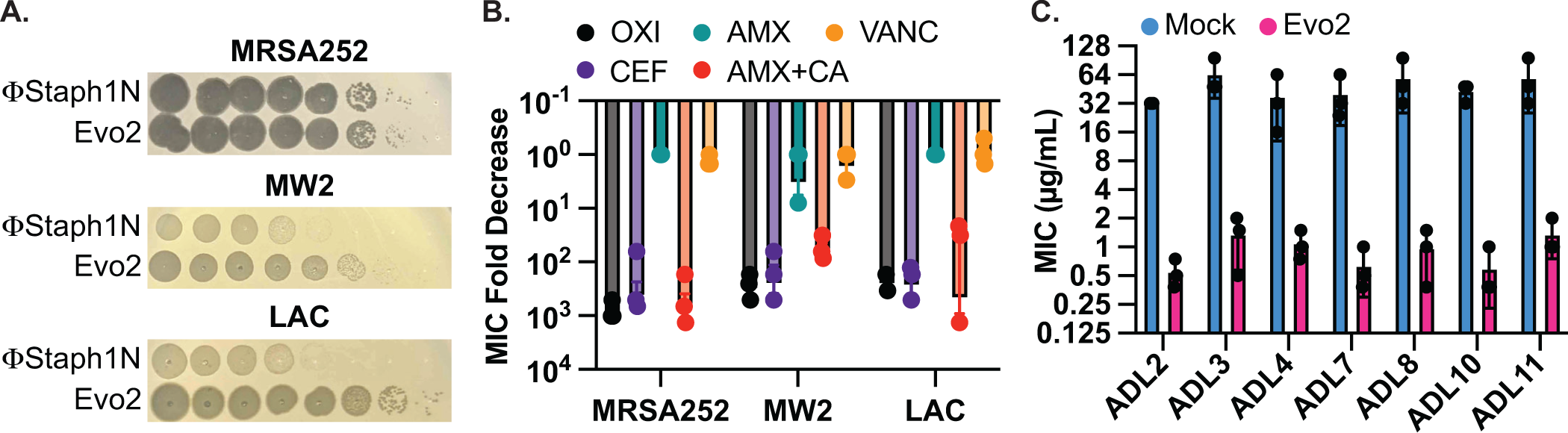
Evo2 is a variant of ΦStaph1N with higher activity against MRSA. **A.** Evo2 shows comparable infectivity towards MRSA252 but improved infectivity towards MW2 and LAC, relative to ΦStaph1N. The same plaquing data is also shown in Figure S3A. **B.** Similar to ΦStaph1N, Evo2 infection reduces β-lactam resistance in MRSA. **C.** Evo2 infection reduces the MIC against oxacillin in clinical isolates of USA300 (ADLs).

We sequenced the genome of Evo2 to determine the genetic mechanism driving this enhanced activity. We observed a single point mutation in ORF141 that induces a premature stop codon (Figure S3B). Sequence analysis with HHpred predicts ORF141 to be a putative DNA binding protein with an HTH motif (PDB: 2LVS, E-value: 2.5e-9). We speculate that this protein is a transcriptional regulator that when inactivated by a nonsense mutation, increases ΦStaph1N infectivity. Future studies will center on determining the mechanism of this mutation and why Evo2 only evolved in the LAC strain.

Given Evo2’s enhanced activity against MRSA, we asked how predation by Evo2 affected β-lactam resistance. We infected MRSA252, MW2, and LAC with Evo2 at an MOI of 0.1 and measured the MICs against β-lactams after 48 hours of passaging. Similar to ΦStaph1N, infection by Evo2 reduced the MICs of the three MRSA strains against OXA, CEF, and AMX+CA, while MICs against AMX alone and VAN did not change significantly (Figure 2B). Expanding on the different classes of antibiotics, we tested whether Evo2 predation could impact the susceptibility to the transcription inhibitor rifampicin; the translation inhibitors erythromycin and mupirocin; and the cell envelope disruptors teicoplanin, fosfomycin, and daptomycin (Table S3). We found that the MICs of these antibiotics did not change significantly, with a few exceptions: in some cases, Evo2-resistant LAC became sensitized to fosfomycin and daptomycin; furthermore, one replicate of Evo2-resistant MRSA252 evolved sensitivity to teicoplanin. However, overall the MIC reduction in these cases was not as dramatic as the MIC reduction seen against β-lactams.

Finally, we examined how different MOIs of Evo2 impacted β-lactam resistance in MRSA (Table S2). We infected the three MRSA strains with Evo2 at varying MOIs from 10^−2^ to 10^−5^ and measured the MIC against oxacillin of the evolved MRSA. Across all three strains, we found that replicate cultures across different MOIs were unable to recover growth following Evo2 infection (Table S2). However, cultures of MRSA252, MW2 and LAC that did regrow displayed a loss of oxacillin resistance, between 10- to 1000-fold. Thus, overall Evo2 displayed a higher infectivity against the MRSA and a greater potency in reducing β-lactam resistance.

### Evo2 is broadly active against recent clinical isolates of *S. aureus* USA300

MRSA252, MW2, and LAC were isolated in 1997, 1998, and in the early 2000s, respectively. We therefore tested if Evo2 can infect more recent *S. aureus* clinical isolates. We compared the plaquing efficiency of Evo2 and ΦStaph1N against 30 USA300 strains that were isolated between 2008 and 2011 at St. Louis Children’s Hospital (Table S1).^26^ We observe dramatic variation in the plaquing efficiency of ΦStaph1N and the 30 strains, while Evo2 exhibited a higher plaquing efficiency in the majority of the 30 strains (Table S4). We then tested how infection of Evo2 impacted OXA resistance in 12 (ADL1-12) of these clinical isolates. We infected the strains with Evo2 at an MOI of 0.1 and, if a phage-resistant population was recovered, measured the OXA MIC after 48 hours of passaging. Interestingly, after 15 independent challenges with Evo2, we were not able to recover phage-resistant populations from ADL1, 5, 6, and 12. This suggests that Evo2 resistance acquisition is a rare event in these strains. In the rest of the strains, we observed that similarly to MRSA252, MW2, and LAC, the OXA MIC was reduced between 10-to 100-fold after Evo2 infection (Figure 2C). Overall, these results highlight the broader host range and activity of Evo2.

### Effect on β-lactam MIC by other phages

We asked whether two additional phages from our collection could elicit MIC reduction against oxacillin in MRSA. The MRSA LAC strain is sensitive to infection by ϕNM1γ6, a lytic version of the temperate phage ΦNM1 of the *Dubowvirus* genus, derived from the *S. aureus* Newman strain.^27–29^. In LAC, ΦNM1γ6 displays plaquing comparable to that of ΦStaph1N, while also showing activity against some of the clinical USA300 isolates (Figure S1, Table S4). Therefore, we infected LAC with ΦNM1γ6 at an MOI of 0.1 and measured the MIC against β-lactams of the surviving cells. While the recovered LAC cultures exhibited resistance against ΦNM1γ6, they did not show a reduction in MIC (Figure S4), suggesting that phage resistance caused by ΦNM1γ6 is uncoupled from β-lactam resistance. We also isolated from the environment a second phage of the *Kayvirus* genus, called SATA8505,^30^ and tested its activity against MRSA. SATA8505 is active against MRSA252, MW2, and LAC (Figure S5), and infection of the three strains caused a rise of phage resistance in MRSA252, MW2, and LAC; (Figure S5A). Similar to ΦStaph1N and Evo2, cells resistant to SATA8505 showed a strong loss of oxacillin resistance (Figure S5B). Altogether, our results with these two phages suggest that the ability to reduce β-lactam resistance is not a universal feature across all staphylococcal phages and call for a more comprehensive analysis of staphylococcal phages and their ability to elicit β-lactam trade-offs.

### Genomic mutations in MRSA strains following phage infection

Following our phenotypic analyses, we examined the genomes of the phage-resistant MRSA. We first sequenced the genomes of three clonal isolates (A-C) from each MRSA strain that underwent ΦStaph1N, Evo2, or a mock infection. We observed that each MRSA strain evolved distinct mutation profiles (Figure 3A). Irrespective of the strain and phage treatment, most mutations were predicted to be substitutions, followed by truncations (Figure S6). Cluster of Orthologous Genes (COG) variants associated with transcription, cell wall/membrane/envelope biogenesis, coenzyme transport and metabolism, and defense mechanisms were the most commonly found categories. Mutations in annotated genes that appeared at least twice across the clonal replicates are summarized in Table 1. Information on all detected genetic variants is listed in Supplementary Table 1.

**Figure 3.**
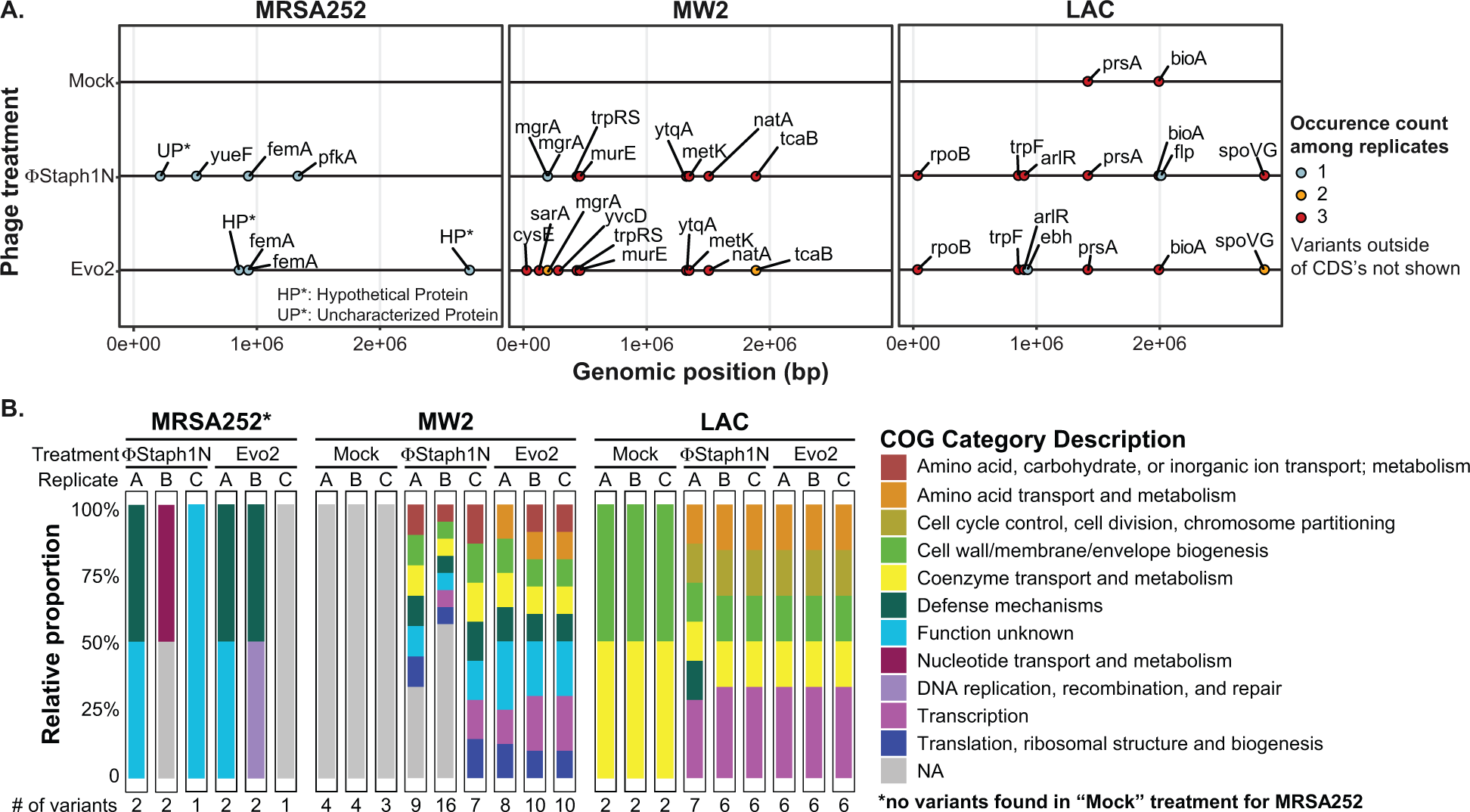
Phage infection of MRSA strains produces distinct mutational profiles. **A.** Coding sequences (CDS) with mutations from the three MRSA strains following phage treatment or mock treatment. For each strain, three isolates were sequenced and their mutations identified. Mutations are color-coded based on the number of occurrences among the three replicates. **B.** Categories of genes with mutations that arose in each MRSA strain and treatment condition.

**Table 1.** Mutated genes in MRSA following infection with phages ΦStaph1N or Evo2.

| Gene | Description | Strain | Phage infection | Reference |
| --- | --- | --- | --- | --- |
| *sarA* | Transcriptional regulator of antibiotic resistance and virulence | MW2 | Evo2 | 31,32 |
| *mgrA* | Transcriptional regulator of antibiotic resistance and virulence | MW2 | ΦStaph1N, Evo2 | 33,34 |
| *rpoB* | Beta subunit of RNA polymerase<br>Transcriptional regulator of antibiotic resistance | LAC | ΦStaph1N, Evo2 | 35 |
| *arlR* | Transcriptional regulator of antibiotic resistance and virulence | LAC | ΦStaph1N, Evo2 | 34,36,37 |
| *spoVG* | Transcriptional regulator of antibiotic resistance and virulence | LAC | ΦStaph1N, Evo2 | 38,39 |
| *cysE* | Cysteine and methionine synthesis, serine O-acetyltransferase | MW2 | Evo2 | 40 |
| *metK* | Cysteine and methionine synthesis, S-adenosylmethionine (SAM) synthetase | MW2 | ΦStaph1N, Evo2 | 41 |
| *trpF* | Phenylalanine, tyrosine and tryptophan synthesis, phosphoribosylanthranilate isomerase | LAC | ΦStaph1N, Evo2 | 42 |
| *femA* | Peptidoglycan synthesis, pentaglycine synthesis | MRSA252 | ΦStaph1N, Evo2 | 43,44 |
| *murE* | Peptidoglycan synthesis, UDP-MurNAc tripeptide synthesis | MW2 | ΦStaph1N, Evo2 | 45 |
| *trpS* | Aminoacyl-tRNA synthesis, tryptophanyl-tRNA synthesis | MW2 | ΦStaph1N, Evo2 | 46 |
| *ytqA* | tRNA modifications, mnm<sup>5</sup>s<sup>2</sup>U synthesis | MW2 | ΦStaph1N, Evo2 | 47 |
| *yvcD* | Unknown | MW2 | Evo2 |  |
| *natA* | ABC transporter | MW2 | ΦStaph1N, Evo2 | 48 |
| *tcaB* | Predicted multidrug efflux pump | MW2 | ΦStaph1N, Evo2 | 49 |
| *fmhC* | Fem-like factors | LAC | ΦNM1γ6 | 50 |

One plausible hypothesis explaining the loss of β-lactam resistance is that phage infection selected for a defective *SCCmec* or *blaZ*. However, we did not observe any mutations in the two loci. Instead, all MRSA strains exhibited mutations in ancillary genes implicated on the loss in β-lactam resistance. In MRSA252, both ΦStaph1N and Evo2 infection were selected for frameshift or nonsense mutations in the *femA* gene that would inactive the protein product. As discussed above, *femA* is required for the synthesis of the pentaglycine branch on *S. aureus* Lipid II, the peptidoglycan precursor (Table 1). Deletions of *femA* have been shown to increase susceptibility to β-lactams even when PBP2a (encoded by *mecA*) is expressed, thus providing a genetic mechanism for how some MRSA252 cells lose β-lactam resistance after phage selection.^43,44^ At the same time, we found the presence of other uncharacterized mutations in phage-resistant MRSA252. For example, clone A of ΦStaph1N-treated cells carried 2 mutations: a frameshift in *femA* and a substitution mutation in an uncharacterized protein; meanwhile, clone B displayed a substitution mutation in *pfkA*, a predicted ATP-dependent 6-phosphofructokinase and mutation in an intergenic region; clone C showed a substitution mutation in a putative transport protein, called *yueF* (Figure 3B, Supplementary Table 1). The role of these mutations in mediating phage resistance or β-lactam sensitivity, if any, remains unknown.

In MW2, we found mutations in two transcriptional regulators, *mgrA* and *sarA* (Figure 3A, Table 1). Both *mgrA* and *sarA* belong to the family of MarR (multiple antibiotic resistance regulator)/SarA (staphylococcal accessory regulator A) proteins, which regulate drug resistance and virulence in *S. aureus*.^31–33^ In ΦStaph1N-treated MW2, only *mgrA* was mutated, while in Evo2-treated MW2, clones also showed nonsense mutations in *sarA.* We also found mutations in *metK* and *ytqA* which are both predicted to be associated with S-adenosylmethione (SAM): *metK* synthesizes SAM, while *ytqA* belongs to the radical SAM enzyme family and is predicted to be involved in tRNA modification.^41,47^ Phage-treated MW2 also displayed mutations in other genes, including *tcaB and murE*. Notably, each clonal replicate had multiple mutations in the genome, while by contrast untreated MW2 cells only displayed a deletion in an intergenic region that is not present in any of the phage-treated samples. These findings suggest that MW2 could be amassing multiple mutations during the course of phage infection.

For LAC, we observed a third, distinct mutational pattern (Figure 3A). Of note, we found nonsense mutations in *arlR,* which is part of the *arlRS* two-component signaling system (Table 1). The activity of *arlRS* has been implicated in *S. aureus* virulence, pathogenicity, and oxacillin resistance.^36,37^ Further, we observed substitution mutations in *spoVG*, which is a transcription factor regulating the expression genes involved in a variety of functions, including cell wall metabolism (Table 1).^38^ Indeed, *spoVG* activates the expression of *femA*.^39^ Studies have shown that *spoVG* modulates β-lactam antibiotic resistance by modulating cell wall synthesis.^39^ Similar to the other two MRSA strains, phage-treated clones of LAC showed multiple mutations in their genomes. We observed mutations in *prsA* and *bioA* that appeared in both the mock and phage treatment conditions, suggesting that these mutations do not arise due to phage selection.

Altogether, our results show that phage-infected MRSA strains acquire distinct mutational profiles. These mutations likely work in concert to promote phage resistance and β-lactam sensitivity, making it challenging to determine the mechanistic contributions of individual mutations. For example, we observed that the genes *mgrA*, *sarR*, and *arlR* evolved nonsense mutations, which would result in truncated, potentially non-functional protein products. We therefore tested if single knockout mutants of these genes alone are sufficient to confer resistance to ΦStaph1N and Evo2 (Figure S7). In MW2, the *mgrA* knockout resulted in a modest reduction in plaquing of Evo2 and ΦStaph1N. However, none of the remaining mutants in either the MW2 or LAC background conferred resistance. Prior experimental studies have also shown that phage resistance in *S. aureus* can arise from the disruption of single genes directly involved in the synthesis and modification of WTA, such as *tagO*.^51^ Some MRSA strains also alter cell wall glycosylation through dedicated genes encoded on prophages.^14^ However, we did not see any mutations in genes directly involved in WTA synthesis. Our results thus highlight how MRSA can take on unique mutational pathways under phage selection.

Finally, we examined the mutational profile in ΦNM1γ6-resistant LAC populations Because infection with ΦNM1γ6 did not result in a decrease in OXA resistance, we hypothesized that mutations that arose in LAC following ΦNM1γ6 would be distinct from those following Evo2 infection. We found that the two genes were mutated across three clonal isolates from different resistant populations: *bioA* and *fmhC.* As seen previously, mutations in *bioA* appeared in the mock treatment, suggesting that that the mutations arose independently of phage selection. On the other hand, LAC showed a missense mutation in *fmhC* (H21D, Table 1, Supplementary Table 1). *FmhC* and its homologue *fmhA* pair with *femA* and *femB* to incorporate Gly-Ser dipeptides into peptidoglycan cross-bridges.^50^ However, the mechanism of the H21D mutation is unknown, and to our knowledge, mutations in *fmhC* have not been associated with phages resistance in *S. aureus* before.

### Phage-treated MRSA strains display broad changes to their transcriptomes

Our genomic analysis revealed that phage-treated MRSA evolved mutations in a variety of transcriptional regulators, some of which are known to affect MRSA virulence. We therefore hypothesized that the mutations in these regulators would fundamentally alter the transcriptional profile of the treated MRSA. To test this, we performed bulk RNA-seq experiments on MW2 and LAC strains that were treated with the phage Evo2 and compared their transcription profiles to those of untreated strains (Figure 4, Table S5). We observed significant changes in gene expression in both MW2 and LAC. Notably, mirroring the trend seen in the mutational data, we did not observe significant changes in the expression of genes in the *SCCmec* cassette or *blaZ* present in both MW2 and LAC.

**Figure 4.**
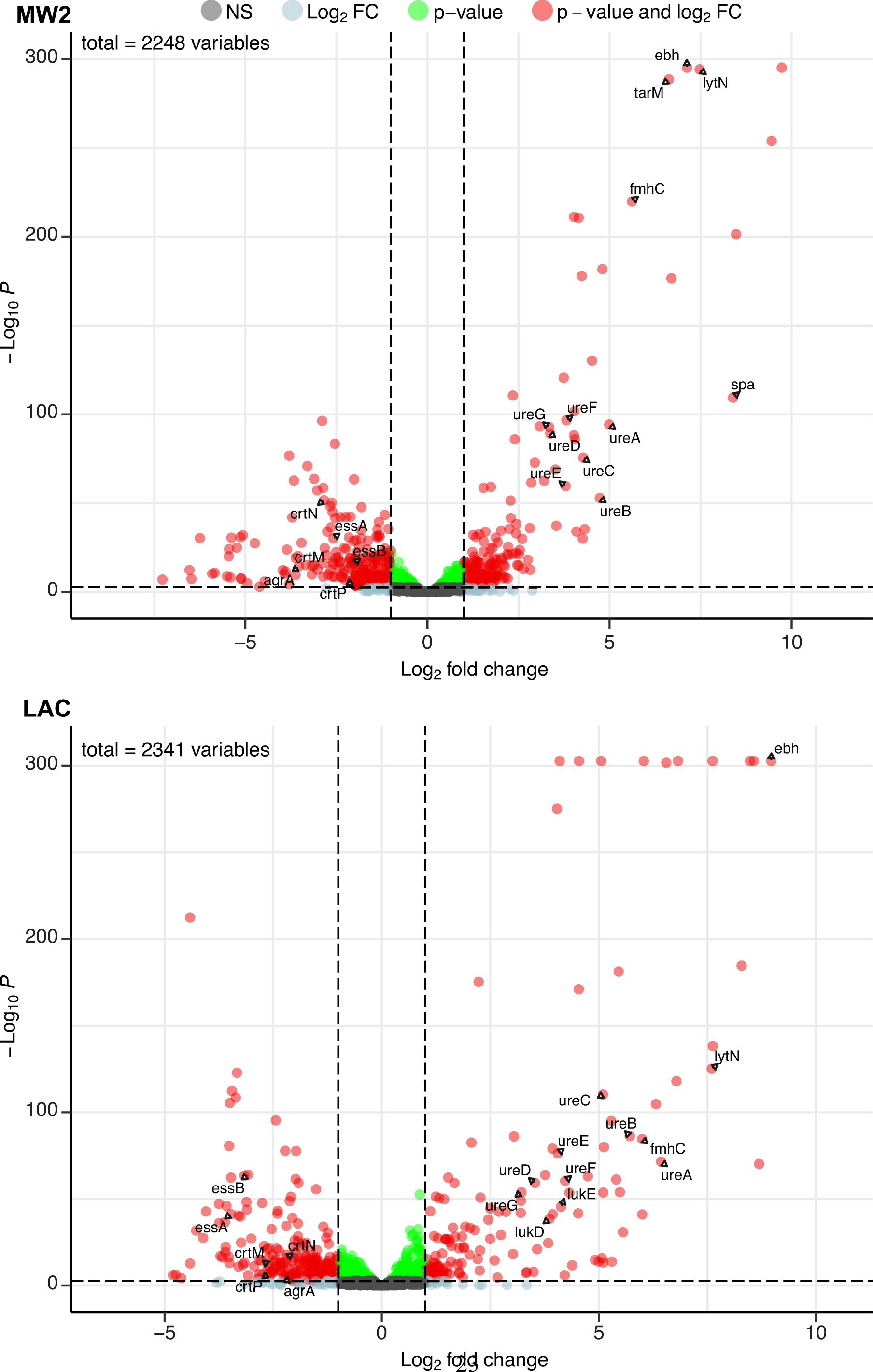
Phage infection changes the transcriptomic profile of MRSA. Differential expression analysis was performed on the transcriptomes of MW2 (top panel) and LAC (bottom panel). For both strains, Evo2-infected samples were compared to uninfected controls. 3 biological replicates were analyzed for each condition. Horizontal dotted lines represent an adjusted p-value cut-off of 0.002, while vertical dotted lines represent a log_2_ fold change of −2 or 2 in expression. Transcripts with a log_2_ fold change between −2 or 2 and a p-value >0.002 are labeled as grey dots (Not significant, NS); transcripts that pass either the fold change or p-value cutoff (but not the other) are represented as blue and green dots, respectively; transcripts that pass both cutoffs are shown as red dots. Genes discussed in the main text are labeled. Data for all the transcripts with significant fold changes is shown in Table S5.

We first compared our expression data against transcriptomic studies from previous studies. For example, Horswill and colleagues have shown that deletions of *arlRS* and *mgrA* de-represses the extracellular matrix binding protein *ebh*, resulting in significantly higher expression levels of the gene. In addition, these deletions also increased the expression of urease genes involved in the urea TCA cycle.^33,37^ In our experiments, both MW2 and LAC strains evolved nonsense mutations in *mgrA* and *arlR,* respectively. We therefore hypothesized that these mutations would mimic the effects of gene deletions and likewise yield elevated transcript levels of *ebh* and urease genes. Aligning with our hypothesis, differential expression data from both MW2 and LAC displayed a log_2_ fold changes of >7 for *ebh* and >3 for *ureABCDEFG* genes (Table S5).

Additionally, both MW2 and LAC strongly upregulated several genes involved in cell wall maintenance. These include *lytN*, which is a murein hydrolase involved in the cross-wall compartment of *S. aureus*, and *fmhC*, which, as described previously, incorporate Gly-Ser dipeptides into pentaglycine cross-bridges in the *S. aureus* peptidoglycan cell wall. Overexpression of *lytN* has been shown to damage the cell wall, which in turn is alleviated by overexpression of *fmhC*.^50^ Interestingly, *fmhC* overexpression is linked to increased β-lactam sensitivity and thus may contribute to the loss of β-lactam resistance phenotypes we observed in the MRSA strains.

MW2 and LAC also downregulated numerous genes, many of which are known virulence factors. Both strains reduced transcript levels of genes in the locus of the type VII secretion system (*ess* locus), staphyloxanthin biosynthesis (*crtM, crtN, crtP*), and quorum sensing (e.g. *agrA*) (Figure 4, Table S5). Individually, these pathways have been shown to bolster the ability of *S. aureus* to establish infection and evade the host immune system.^52–54^ Our results suggest that infection by Evo2 can lead MRSA to reduce the expression of all of these pathways, which could reduce the virulence of *S. aureus*.

Finally, we noted that both MW2 and LAC showed transcriptional changes that appear to be strain specific. For example, MW2 saw significant increase in the virulence factor *spa*, known to interfere with the host immune response and interface with other bacterial species. The presence of cell wall-bound Protein A has also been shown to decrease phage absorption, likely by masking WTA.^55^ Further in MW2, we found that *tarM*, which adds α1,4-GlcNAc to WTA, was strongly up-regulated (log_2_ ratio = 6.63). This is in line with previous findings showing that elevated *tarM* and α1,4-GlcNAc-WTA can lead to phage resistance in MRSA. The LAC showed an increase in the expression (log_2_ fold change >3.8) of the hemolytic cytotoxin genes *lukD/E,* which lyses host cells and targets neutrophils.^56^ We do not know whether the increased expression of these genes results in a greater level of protein production and secretion, but these transcriptional changes could represent a potential “trade-up” associated with phage resistance. Additional studies will be needed to fully assess the physiological and ecological effects of these upregulated genes in MRSA. Altogether, our RNA-seq results suggest that phage infection and resistance in MRSA cause significant transcriptional changes across a wide range virulence, metabolic, and cell-wall associated genes.

### Phage-treated MRSA strains display reduced virulence phenotypes

*S. aureus* is a highly virulent pathogen, relying on a vast array of toxins and immune evasion proteins to promote infection.^57^ MW2 and LAC strains as models of MRSA virulence.^33,34^ In light of our mutational and transcriptomic data, we hypothesized that phage-treated MRSA cells would display altered virulence phenotypes, in addition to reduced β-lactam resistance. We first tested the ability of MRSA strains that survived phage predation for their ability to form biofilms in a Crystal Violet assay. We found that Evo2 infection of MRSA252 resulted in a significant reduction in Crystal Violet absorption compared to the parental strain. However, we show no significant difference in Crystal Violet absorption between parental, mock- and phage-treated MW2 and LAC strains (Figure S8).

Next, we tested whether phage infection could affect the hemolysis of rabbit blood cells. Hemolysis is mediated by the secretion of toxins, notably alpha toxin encoded by the gene *hla*, and plays an important role in MRSA infection.^58^ Expression of these toxins is regulated by virulence pathways that comprise numerous transcription factors, including *mgrA*, *arlR*, and *sarA.* Furthermore, in our RNA-seq results, we found that phage-resistant MRSA strains showed reduced expression of other cytotoxins. Parental MW2 and LAC colonies lysed rabbit blood cells on blood agar plates, producing distinct halos of clearance around the bacterial cells. For untreated LAC, the total area of hemolysis was on average 3-fold larger than that of untreated MW2 (∼210 mm^2^ vs ∼70 mm^2^, respectively); with MRSA252, by contrast, lysis was not detected (Figure 5A). Following treatment with ΦStaph1N, we observed that MW2 and LAC displayed a reduced area of hemolysis by 4 to 5-fold. With Evo2-treated cells, we found that in MW2 the fold reduction was comparable to that of ΦStaph1N-treated cells. However, for Evo2-treated LAC, loss of hemolysis was even more pronounced, with two of the replicates showing no detectable hemolysis. We note that neither MW2 nor LAC showed a reduction in transcript expression of *hla.* We posit that the loss of hemolysis could be driven by an inability of phage-evolved MRSA to secrete the toxin.

**Figure 5.**
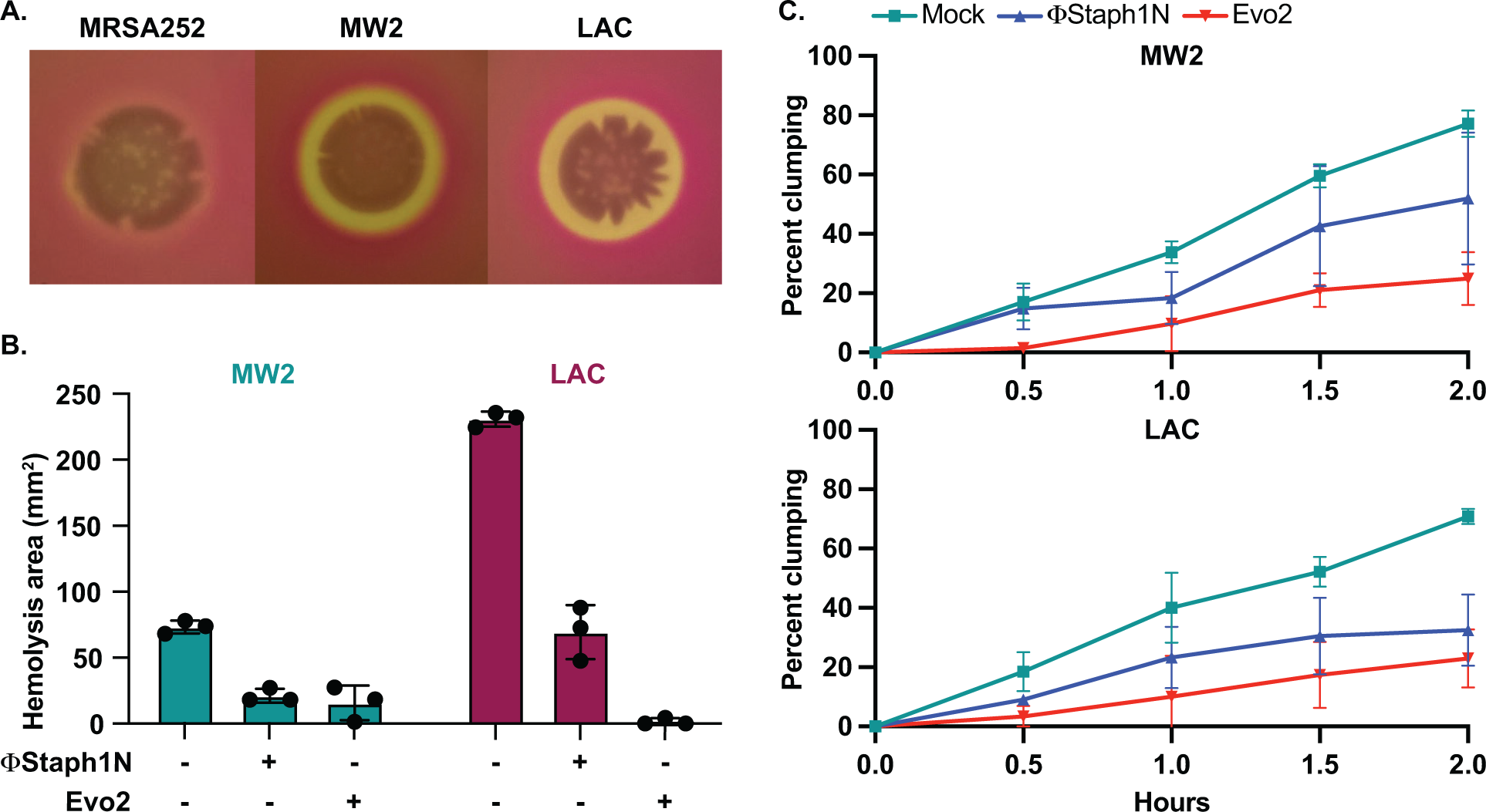
Phage treatment of MRSA results in attenuated virulence phenotypes. **A.** MW2 and LAC strains display hemolytic activity on rabbit blood agar plates, while MRSA252 does not. **B**. Phage-treated MW2 and LAC strains display reduced hemolysis compared to uninfected cells. **C**. Surviving cultures of MW2 and LAC treated with either ΦStaph1N (blue) or Evo2 (red) show reduced clumping rates compared to mock untreated cells (teal).

We next tested how phage infection affected cell agglutination (or clumping) in MRSA. *S. aureus* binds to fibrinogen, forming protective aggregates of bacterial cells. Clumping is thought to have several functions in the context of staphylococcal infections, facilitating adhesion to the host tissue. Clumps are also likely to be more resistant to clearance by the immune system, partly because they may be too large to be phagocytosed by neutrophils.^34^ In our transcriptional data, we noted that several cell surface proteins known to reduce cell clumping, such as *ebh,* were over-expressed in phage-resistant MRSA. Horswill and colleagues found that de-repression of *ebh* reduces clumping. Indeed, phage-treated MW2 and LAC displayed less clumping than the mock-treated or parental strain. For MW2, ΦStaph1N infection resulted in a modest reduction, while Evo2 infection resulted in a reduction of approximately 3-fold (Figure 5B). In LAC, we found that both ΦStaph1N and Evo2 treatment resulted in comparable reductions of clumping in surviving cells. Overall, these phenotypic results align with our genetic and transcriptomic data, showing that phage infection can drive MRSA populations to reduced virulence phenotypes.

### Combination treatment between Evo2 and β-lactam

The aforementioned results suggest that MRSA cells evolve phage resistance following infection, which is associated with trade-offs in virulence and β-lactam resistance. We next asked how MRSA populations would evolve under co-treatment with phage and β-lactam. In principle, these two simultaneous selective pressures could drive the evolution of resistance against both the phage and antibiotic, negating the trade-offs in drug resistance. To test this, we performed checkerboard assays with phage and oxacillin on MRSA252, MW2, and LAC. Serial dilutions of Evo2 or ΦStaph1N were mixed with serial dilutions of oxacillin on a 96-well plate (Figure 6A), after which MRSA strains were added to the plate and allowed to grow for 24 hours. Following 24 hours, one percent of the culture in each well was transferred into a fresh plate well with nonselective media, and the cultures were allowed to grow for another 24 hours (48 hours total). Throughout the experiment, the cell density was monitored by measuring the optical density in each plate well.

**Figure 6.**
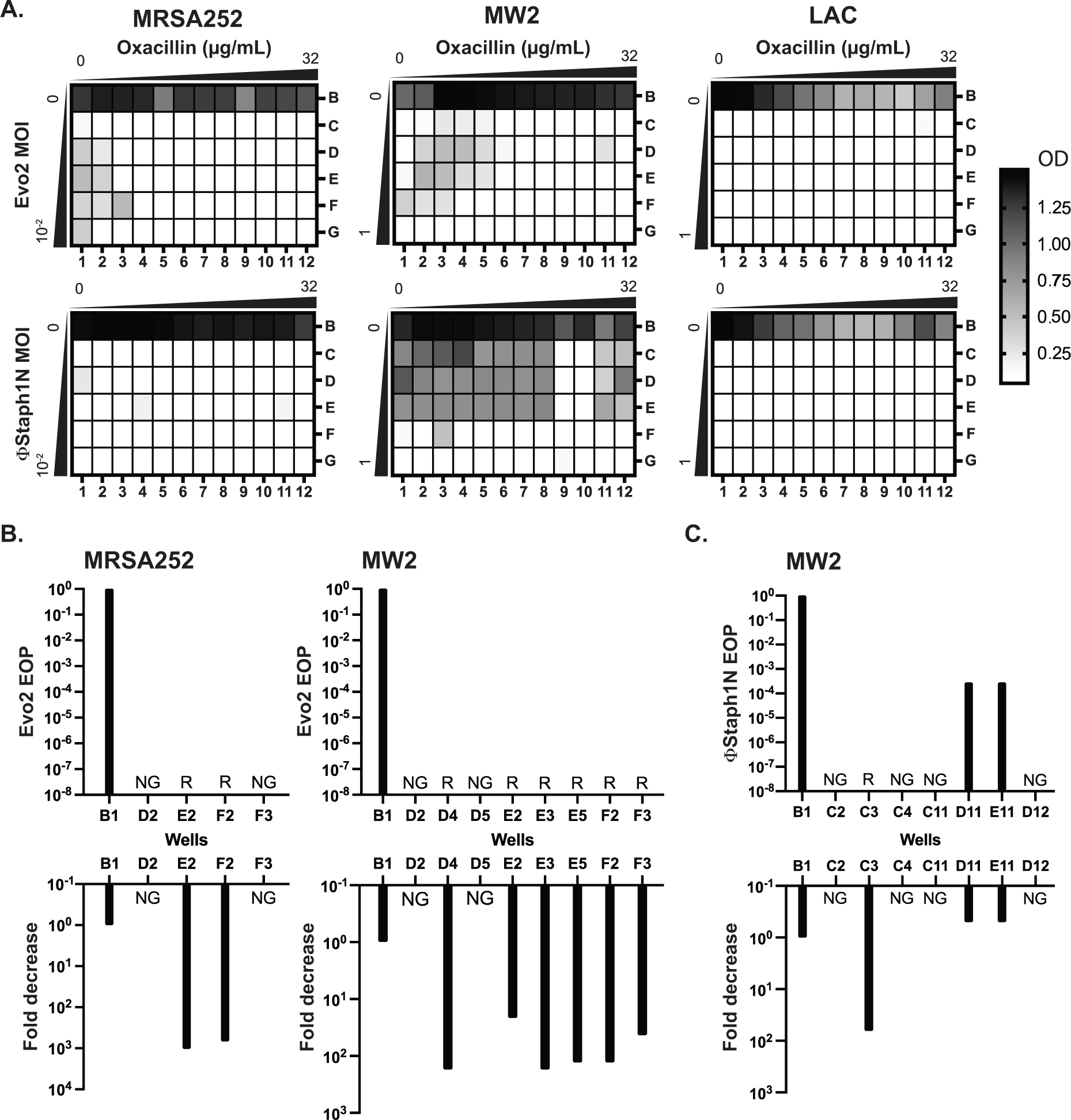
Co-treatment of MRSA with bacteriophage and β-lactam. **A.** Checkerboard assays of MRSA strains with gradients of oxacillin and Evo2 (top panels) or ΦStaph1N (bottom panels). The oxacillin gradient is a 2-fold serial dilution of drug concentration (µg/mL), while the phage MOI gradient is a 10-fold serial dilution of MOI. The rows and columns of each plate are labeled with letters and numbers, respectively. The black-white gradient in each well reflects the optical density of the culture and is the mean value from three biological replicates. MRSA strains co-treated with oxacillin and (**B)** Evo2 or (**C)** ΦStaph1N were tested for their phage resistance and oxacillin resistance. The letter/number combination reflects the well from which the cells were picked for analysis. Wells that could not produce a viable culture are labeled as NG (no growth). For wells that regrew, we calculated the efficiency of plaquing (EOP) of phage and measured the fold reduction in oxacillin MIC. Cultures that showed no detectable viral plaques are labeled as resistant (R).

We first examined how MRSA grew in combinations of Evo2 and antibiotic (Figure 6A, top row). For MRSA252, cells grew at low levels of phage (MOI of 0.01 or less) and oxacillin (<0.125 µg/mL) (Figure 6A). LAC displayed greater sensitivity, displaying no detectable growth after 48 hours in the presence of phage, irrespective of the presence of oxacillin. For MW2, cells showed limited growth at MOIs <1 and oxacillin levels <0.125 µg/mL. For each strain, we picked cells from wells with an OD_600_ >0.5 and inoculated them into fresh, non-selective media. As a control, we also regrew cells that were treated with neither phage nor oxacillin (well B1). For MRSA252, two (E2 and F2) wells contained viable MRSA cells, forming turbid cultures; for MW2, six (D4, E2, E3, E5, F2, and F3) wells picked regrew (Figure 6B). We posit that cells in the failed cultures had reduced viability from the phage and antibiotic treatment. We tested these regrown cultures for their phage and oxacillin susceptibility. As expected, MRSA252 and MW2 cells from the B1 control wells were sensitive to Evo2, but resistant to oxacillin. In MRSA252, cells from E2 and F2 were resistant to Evo2 infection and exhibited a 1000-fold decrease in MIC against oxacillin. Similarly, in MW2, the six viable cultures exhibited Evo2 resistance and a 10- to 100-fold decrease in MIC. Altogether, these results match with those of MRSA that underwent single selection with Evo2.

We next analyzed how MRSA grew under ΦStaph1N/β-lactam combinations (Figure 6A, bottom row). Neither MRSA252 nor LAC could grow in any ΦStaph1N/oxacillin combination; MW2, by contrast, grew across under a wide range of ΦStaph1N/oxacillin combinations. When ΦStaph1N-infected MW2 was treated with high (16 µg/mL) levels of oxacillin, recovered cells (wells D11 and E11) showed no reduction in MIC against oxacillin (Figure 6C). However, the plaquing efficiency of ΦStaph1N was reduced by 3 to 6 orders of magnitude. Whole genome sequencing revealed that these cells evolved a unique set of mutations different from those seen in the single phage treatment conditions (Table S6). However, when ΦStaph1N-infected MW2 was co-treated with low levels (<0.125 µg/mL) of oxacillin, recovered cells (C3) displayed strong phage resistance and a 100-fold reduction in oxacillin resistance, mirroring phenotypes observed in the single phage treatment experiments.

We note that of the three MRSA strains tested, MW2 is the least sensitive to ΦStaph1N infection. We posit that the selective pressure exerted by high levels of β-lactam dominates over the selective pressure of ΦStaph1N, leading to the evolution of cells with continued β-lactam resistance and limited phage resistance. However, at lower β-lactam levels, the pressure exerted by phage infection predominates, leading to the rise of cells with complete phage resistance and trade-off in β-lactam resistance. This dose-dependent selection by oxacillin is not observed when a more active phage (e.g. Evo2) is used. While limited to one MRSA strain, these results suggest that different phage/β-lactam combinations can produce divergent evolutionary outcomes in MRSA, each with potential clinical implications.

## Discussion

Here we report the discovery that infection by certain phages can drive MRSA populations to evolve favorable genetic trade-offs between phage and β-lactam resistance. Exploiting genetic trade-offs has been proposed as a means to combat resistance in bacterial pathogens.^18^ Not only could phage treatments reduce the bacterial load of an infection, but also potentially resensitize bacterial populations to antibiotics against which they were previously resistant. We show that MRSA strains infected with *Kayviruses* ΦStaph1N, Evo2, and SATA8505 evolved resistance against phage, yet developed up to a 1000-fold loss in their MICs against β-lactam antibiotics. In addition, these evolved MRSA display reduced virulence phenotypes such as lower levels of hemolysis and clumping. Our findings show that phages can resensitize MRSA to β-lactams and even decrease their virulence, which are outcomes of significant biomedical value. However, not all staphylococcal phages can mediate these trade-offs: infection with ΦNM1γ6, a phage of the *Dubowvirus* genus, did generate phage resistance in MRSA, but did not produce a drop in β-lactam resistance. A major future direction will be to determine which types of phages elicit these beneficial evolutionary trade-offs in MRSA.

Our results also paint a complex picture of MRSA evolution during phage infection. Whole genome sequencing revealed that MRSA strains evolved distinct mutation profiles following phage infection, suggesting a multitude of evolutionary paths that different bacterial strains can undertake to evolve resistance against phage. Not only did MRSA strains evolve distinct mutations, but individual, phage-resistant clones accumulated multiple mutations in their genome. Nonetheless, different MRSA displayed a convergence of phenotypes in the form of phage resistance, reduced β-lactam resistance, and attenuated virulence. We posit that this convergence is caused by the involvement of the cell wall in all three phenotypic outcomes. Phages must interface with the cell wall, β-lactams target proteins associated with cell wall maintenance, and many *S. aureus* virulence factors are embedded within the cell wall or are secreted through it. Thus, any modifications to the integrity or chemical composition of the cell wall by phage resistance will impact β-lactam sensitivity and virulence. Cell wall maintenance is controlled by numerous genes, ranging from single proteins involved in cell wall synthesis, such as *femA*, to transcriptional regulators, such as *mgrA*, that control the expression of cell wall synthesis genes. Thus, the mutational patterns observed in each MRSA strain could reflect genetic solutions that enable the bacteria to adapt to the phage predation, while also maximizing the fitness for that particular strain.

Strikingly, MRSA heavily modulated transcriptional profiles following phage infection. We believe these altered expression profiles are a consequence of the genomic mutations that emerged in the various transcriptional regulators. We observed that evolved cells downregulated genes involved in quorum sensing, type VII secretion, and a variety of toxins. It is intriguing to speculate how the down-regulation of these genes impacts MRSA interactions with other bacteria occupying the same ecological niche and with the host immune system. At the same time, MRSA strains also upregulated expression of select virulence factors, such as *spA*, which could represent “trade-ups.” Trade-ups are thought of as non-selected traits that are enhanced following selection (e.g. cross-resistance between phage and antibiotic), which from a therapeutic perspective might be undesirable.^59^ Future work will focus on assessing the risk of these trade-ups in light of the clinical benefit of reduced resistance and virulence.

Drug resistance in bacterial pathogens is an evolutionary problem and will require evolution-guided solutions to mitigate. Our findings highlight the ability of phages to dramatically alter the evolution and physiology of drug-resistant MRSA. Select phage treatments can force bacterial populations down evolutionary paths that make them vulnerable to antibiotics or the host immune system. Critically, this permits the re-deployment of agents that would otherwise remain ineffective, buying time for new drug discoveries. We therefore hope that our work may suggest avenues of research into new phage-based treatment strategies against MRSA and other drug resistance pathogens.

## Acknowledgments

MT is supported by the SciMed GRS program at UW-Madison; AJHV is supported by the NSF Graduate Research Fellowship Program; CYM is supported by start-up funds from the Department of Bacteriology at UW-Madison and the Margeret Q. Landenberger Foundation. We are grateful to Dr. Wilmara Salgado-Pabón, Dr. Petra Levin, and Dr. Alexander Horswill for providing us with MRSA strains and helpful suggestions. We thank the Marraffini and Hatoum-Aslan laboratories for providing us with bacteriophage samples. Computational analyses were performed using the UW-Madison Center for High Throughput Computing. We also thank all members of the Mo lab for their scientific input.

## Author contributions

MT and AJHV contributed equally to the work. MT, AJHV, and CYM designed the study; MT and AJHV performed all experiments; PQT performed the bioinformatic analysis; EDL and LT isolated and tested the bacteriophage SATA8505; CYM supervised the project. MT, AJHV, and CYM wrote the manuscript, with input from the other authors.

## Online Methods

### Strains and culture conditions

The bacterial strains used in this study are listed in Table S1. Unless otherwise indicated, all MRSA strains were grown in Brain Heart Infusion (BHI) media at 37 °C with shaking (235 RPM).

### Plate-based plaque assay

Bacterial lawns were prepared by mixing 100 µL of an overnight culture with 5 mL of melted BHI agarose (top agar). The bacteria and top agar mixture were poured onto a solid BHI plate. The plate was dried for 10 minutes. 10-fold serial dilutions (10^0^ −10^−7^ unless otherwise noted) of phage were then spotted on the bacterial lawn. Plates were then incubated at 37 °C for 16 hours. Phage titer in plaque-forming units per µL (pfu/µL) was then calculated.

### Phage infection assay

MRSA strains were plated onto BHI agar plates and grown overnight. Individual colonies from the parental strains (also referred to as P0) were picked. Single colonies were inoculated in a round bottle tube containing 5 mL BHI broth. The cultures were incubated at 37 °C, 235 RPM for 24 hours. The grown P0 cultures were then diluted 1:100 into fresh 5 mL BHI broth. At the early log phase (OD ∼0.3), the bacterial cultures were treated with phage at an MOI of 0.1, unless indicated otherwise. The treated bacterial cultures were incubated at 37 °C with shaking (235 RPM) for 24 hours. The cultures were then passaged 1:100 into fresh 5 mL BHI broth. This passage was then grown at 37 °C, 235 RPM, for another 24 hours. Surviving cultures were then used for both phenotypic assays and sequencing experiments. As a negative control, MRSA strains were passaged using the steps described above without phage treatment (mock).

### MIC assay

Bacterial lawns were normalized to contain 1×10^8^ CFU/mL bacteria mixed with top agar for a total volume of 5 mL. The bacteria and top agar mixture is poured onto a solid BHI plate. The plate was dried for 10 minutes. MIC with increasing concentrations of antibiotics were placed on the semi-dried bacterial lawn and allowed to dry for 10 minutes. The plates are then incubated at 37 °C overnight. For analysis, the plates were imaged, and the MIC of the bacteria was determined. The MIC is determined at the edge of the inhibition ellipse intersects the side of the strip.

### Rabbit blood hemolysis

Phage-treated or mock-treated cultures were diluted to an OD600 of 0.1, 5 µL of this dilution was spotted on rabbit blood TSB agar plates and incubated at 37 °C for 24 hours. The area of clearance was determined by the following formula: [π (diameter of clearance/2)^2^] − [π(diameter of bacterial spot/2)^2^]

### Clumping assay

Clumping assays were performed as described previously.^60^ In short, overnight cultures were diluted 1:100 and incubated at 37 ℃ until the cultures reached an OD_600_ of 1.5. At this point, 1.5 mL of culture was washed two times and resuspended with PBS. Lyophilized human plasma was added for a final concentration of 1.25%. Resuspended cells were left to sit statically at room temperature. 100uL were taken from the top of the cell suspensions in 30-minute intervals and the OD_600_ was measured.

### Biofilm assay

Biofilm assay was performed using the crystal violet method as outlined.^61^ In brief, overnight cultures grown in BHI at 37 °C were back diluted 1:100 into a 96-well bottomed microwell plate. The plates were incubated without shaking at 37 °C overnight. The contents in the plate were discarded and washed with PBS. Biofilm fixation was done with sodium acetate (2%). Crystal violet (0.1%) was used for staining followed by a final wash with PBS. Absorbance at 600 nm was read using a spectrophotometer.

### DNA sequencing and genome assembly

Following published protocols, genomic DNA from bacteria and phage was isolated using phenol-chloroform extraction. Purified DNA was sent to Plasmidsaurus and SeqCenter for Nanopore and Illumina sequencing, respectively. Reference genomes for bacterial strains were assembled using Flye v2.9.3 with default settings for long-reads. This resulted in 1 singular contig assemblies for 252 (2902592bp, 125x) and MW2 (2820460, 600x), and 3 contigs for LAC (2907712, 645x). Phage assemblies for Evo2 and ΦStaph1N were done with the SPADES assembler v3.15.5.

Open-reading frames (ORFs) were called on the assembled bacterial genomes using Prodigal v.2.6.3,^62^ resulting in a gene-feature file (GFF), and translated genes as .faa and .fna formats. We used BLASTp (Accessed June 4th, 2024), against the protein BLAST database swissprot_2023-06-28, with an expectation value cutoff of 0.001. The top hit for each ORF was used as the final functional annotation. Additionally, we annotate the genomes using Bakta v.1.10.3.

### Mutation identification

A total of 27 genomic samples were collected. DNA was extracted and sent for long-read sequencing using Oxford Nanopore Technology (Plasmidsaurus, San Francisco, USA). Reads were filtered using filtlong v0.2.1 using default settings (https://github.com/rrwick/Filtlong) and with the −p flag 95 (keeping 95% of the best reads). Quality-filtered long reads were mapped against the respective genomes using minimap2 2.22-r1101^63^ resulting in 1 alignment file output per sample (.sam file). The read mapping software Minimap2^63^ was selected because of its suitability to map long reads. Samtools v1.20^64^ was used to convert the .sam files into .bam files, sort the bam file, index the bam files, and generate a coverage table for each position along the alignment.

Reference files (fasta and GFF files) and the 53 “sorted.bam” alignments files were imported into Geneious. Variant calling was performed using Geneious Prime Geneious Prime ® 2024.0.2 (www.geneious.com), using the custom settings: 10% coverage, minimum 95% variant frequency threshold, and the option for “Analyze effect of variants on translation” checked. Variant results and genome annotations table files were exported as a tab-separated-table, and visualized using R v4.4.0, mostly with the tidyverse package. Sequencing data processing, quality filtering, and mapping were performed at the Center for High-Throughput Computing (https://chtc.cs.wisc.edu/). BLAStp was performed on usegalaxy.eu (Accessed June 4^th^, 2024).

### RNA purification

Parental and evolved MRSA strains were diluted 1:100 in BHI broth and incubated at 37 °C, 235 RPM until they reached an OD_600_ of 1.5. 500 µL of culture were transferred into a microcentrifuge and 1 mL of RNAprotect Bacteria reagent (QIAGEN) was added. The mixture was vortexed for 5 seconds and incubated at room temperature for 5 minutes. The tubes were centrifuged for 10 minutes at 5000g, and the supernatant discarded. Bacterial pellets were resuspended in 80 µL of phosphate buffered saline and 10 µL of lysostaphin solution (1 mg/mL stock). The suspensions were incubated at 37 °C, with shaking, until the solution looked clear (∼30 minutes). 10 µL of 10% sarkosyl was then added and the tube mixed, after which 300 µL of TRIzol reagent (Invitrogen) was added. RNA purification was performed following the protocol from the Direct-zol RNA Miniprep Plus kit (ZYMO Research). DNase I treatment was performed as recommended, and the samples were eluted in 75 µL of DNase/RNase-free water.

### RNA sequencing and differential gene expression analysis

Purified RNA was prepared and sequenced on an Illumina sequencing platform at the UW-Madison Gene Expression Center. RNA-seq data was collected in the parental and evolved MRSA strains to assess differentially expressed genes. Cleaned RNA reads were mapped onto the LAC and MW2 reference genomes using bowtie2 v2.5.4,^65^ and featureCount v2.0.8^66^ from the software subreads was used to generate a read count matrix. Two read count matrices (one for LAC and one for MW2) were imported into R v.4.4.0 for processing with DESeq2 (version 1.44.0).^67^ Figures were generated using the packages tidyverse (version 2.0.0) and EnhancedVolcanoPlots (version 1.22.0). To generate the volcano plots, we chose an adjusted p-value of 0.002 and a log_2_ fold change (log_2_FC) of < −2 or > 2. Multiple “unknown” genes were deemed significant (adjusted p-value < 0.002, abs(log_2_FC) >= 2) in both LAC and MW2. To compare the results between the genomes, we performed a protein clustering analysis using MMseqs2 version b804fbe384e6f6c9fe96322ec0e92d48bccd0a42 between all the Bakta-generated amino acid sequences (.faa files) from LF and MW2.^68^ Then we considered any protein sharing over 80% identity to be the “same” to generate a summary table showing up-regulation and down-regulation among the 2 genomes.

### Checkerboard assay for phage-antibiotic synergy

The overnight cultures of MRSA252, Lac-Fitz, and MW2 were back-diluted 1:100 in BHI and incubated at 37 °C until the culture reached mid-log phase. The culture was then inoculated into each well of the 96-well plate containing a gradient of oxacillin and phage (∏Staph1N or Evo2). The oxacillin gradient was a 2-fold serial dilution, while the phage MOI gradient was a 10-fold serial dilution. The plates were then placed at 37 °C with shaking (235 RPM) for 24 hours.

Following 24 hours, each well from the plate was then passaged 1:100 into another 96-well plate with fresh BHI and grown at 37 °C with shaking (235 RPM) for another 24 hours. Following 48 hours of growth, the OD_600_ was measured with a plate reader. Each checkerboard assay was performed in three biological replicates. A selection of surviving cells from individual wells (cells with an OD_600_ reading of > .5) were then picked for phage plaquing and MIC assays. Efficiency of plaquing (EOP) was calculated by the following formula: plaque titer of treated cells/plaque titer of non-treated cells. Determine the mutation profile of survivor cells, genomic extraction, Illumina sequencing, and mutational analysis described above were used.

## Data and code availability

All the scripts can be found at: https://github.com/patriciatran/mrsa-project

## Supplemental data

**Figure S1.**
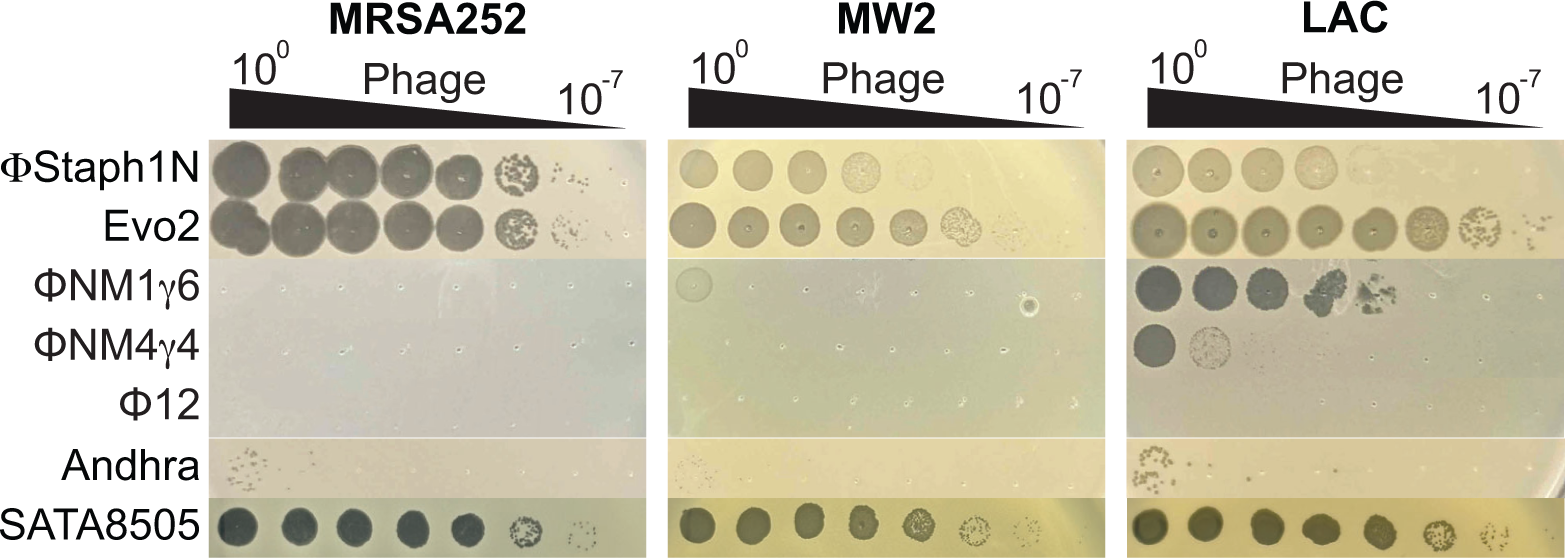
Phage sensitivity of MRSA strains. Efficiencies of plaquing of phages on MRSA252, MW2, and LAC. Phages were 10-fold serially diluted and spotted onto top agar overlays of each strain.

**Figure S2.**
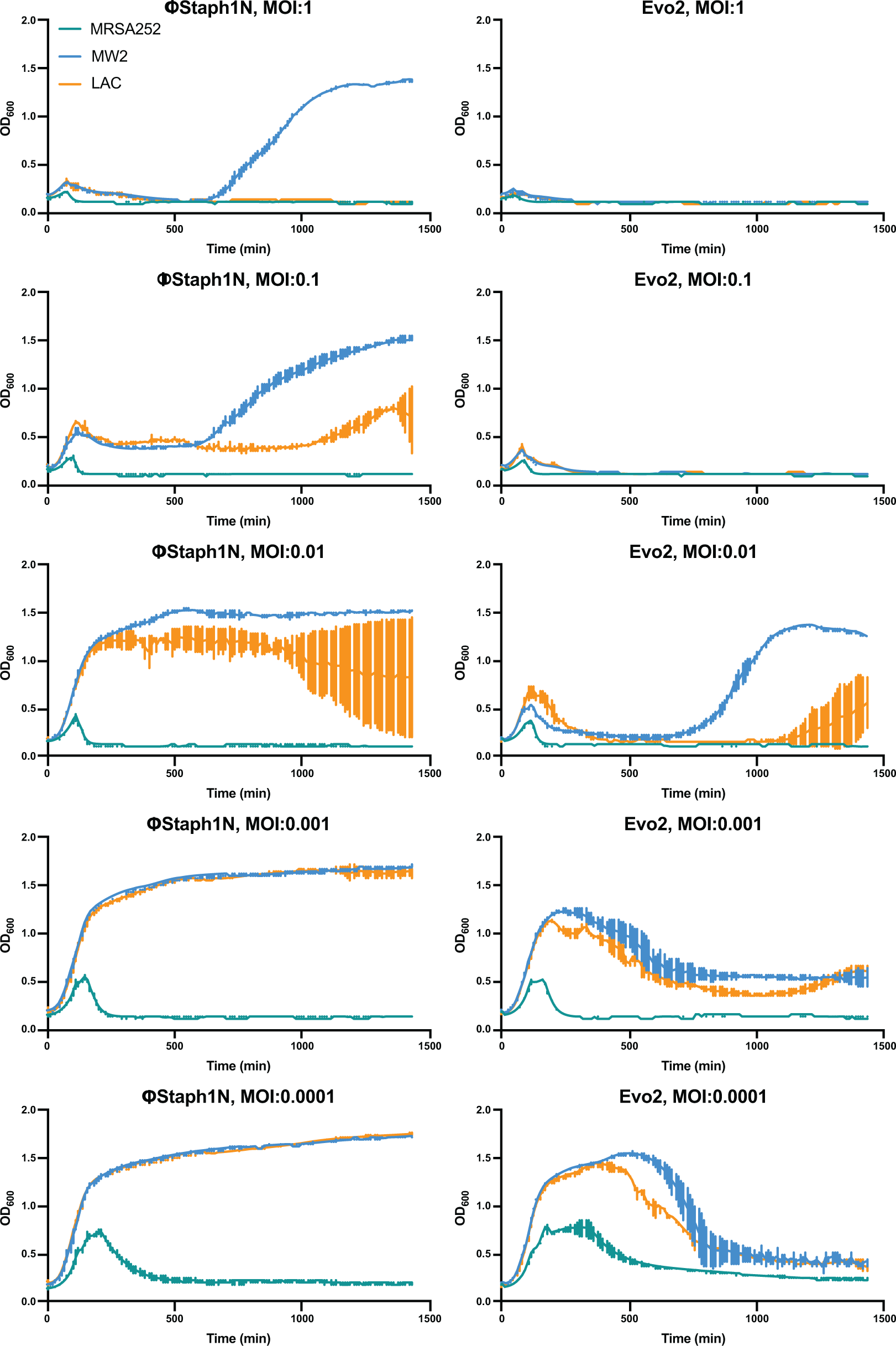
Growth curves of MRSA strains under varying levels of phage infection. MRSA252, MW2, and LAC cultures were infected with either ΦStaph1N or Evo2 at the indicated multiplicity of infection (MOI). The optical density (OD_600_) of the cultures was monitored on an automated plate reader. Each condition was tested in 3 independent replicates.

**Figure S3.**
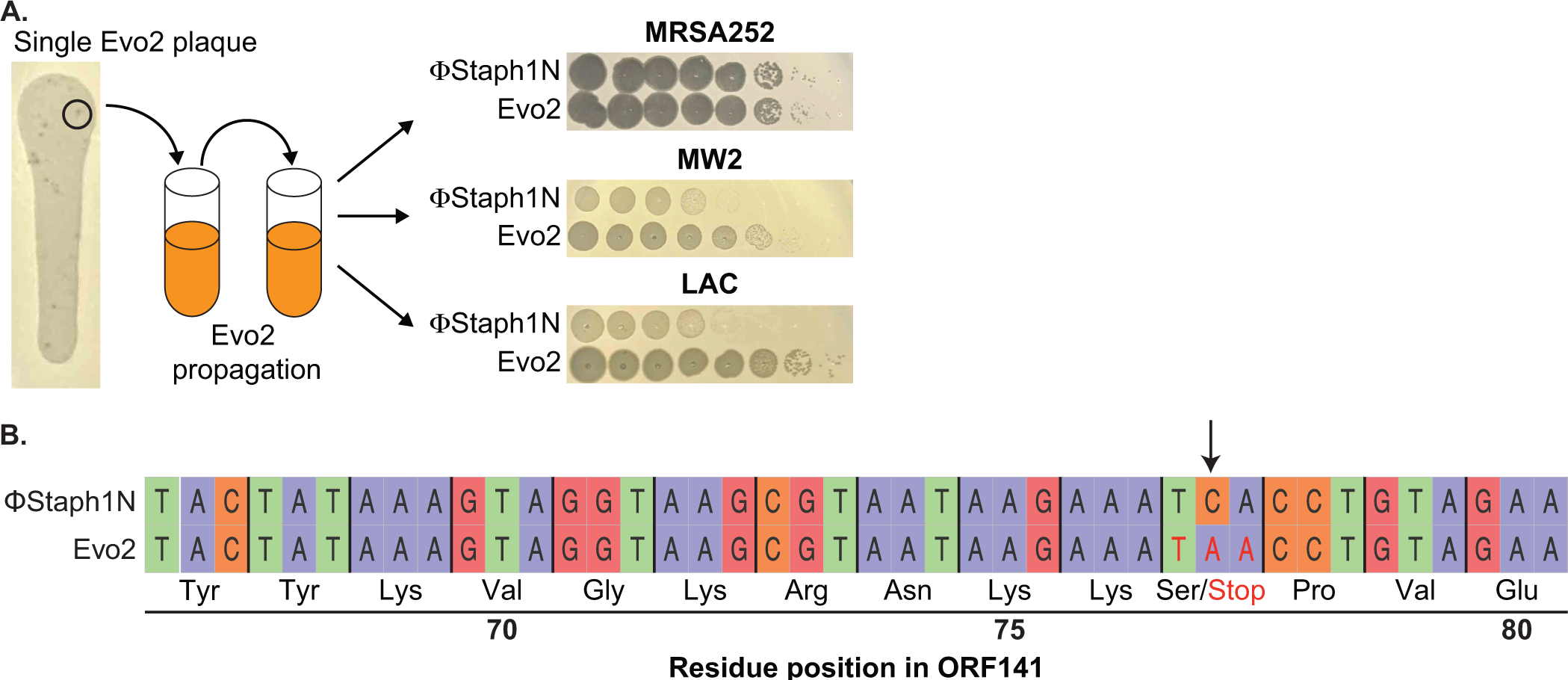
Isolation and sequencing analysis of Evo2. **A.** Individual Evo2 plaques appeared in larger ΦStaph1N plaques on LAC. Individual plaques were isolated and propagated in liquid culture. Evo2 shows improved plaquing on MW2 and LAC. Plaquing data in the right panels are the same as in Figure 2A. **B.** Evo2 is a mutant form of ΦStaph1N with a nonsense mutation in ORF141. The A to C mutation (marked by the arrow) in Evo2 converts Serine 77 of ORF141 into a stop codon.

**Figure S4.**
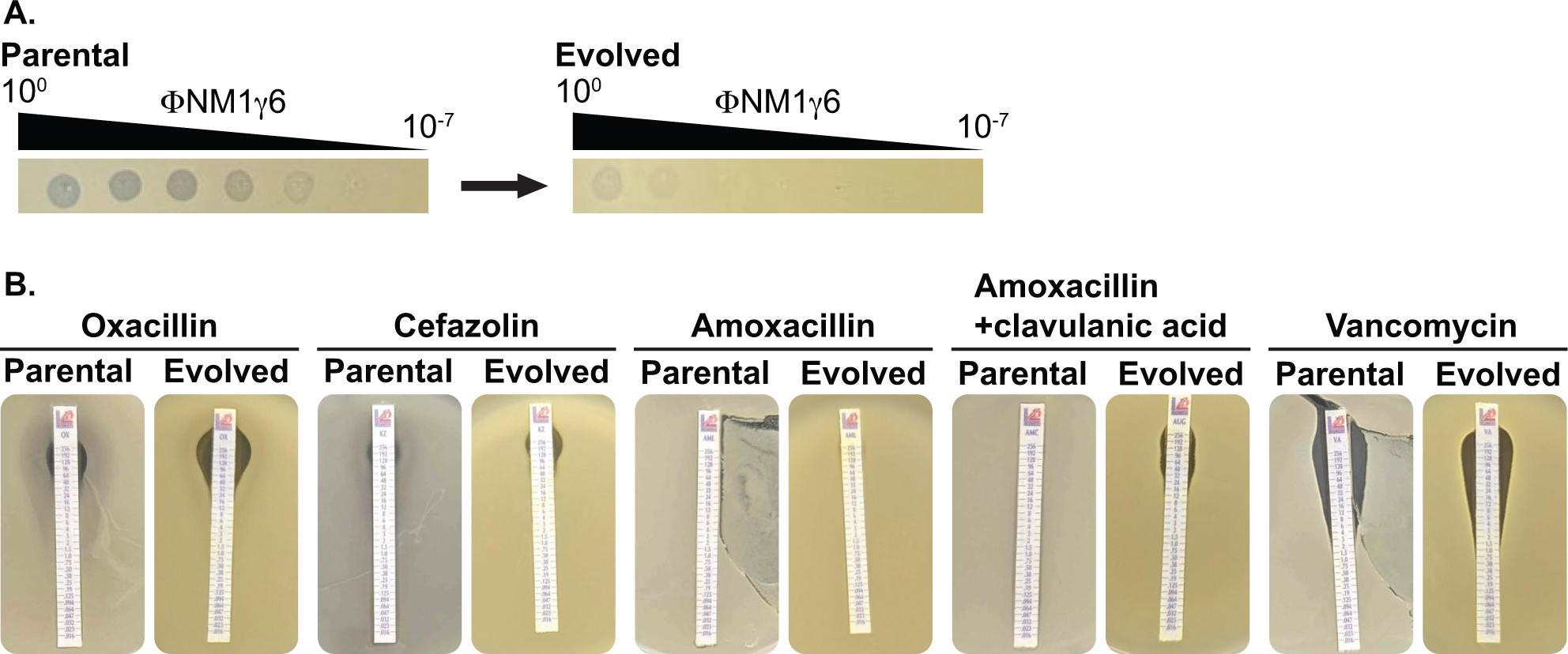
Phage ∏NM1γ6 infection LAC does not drive the loss of β-lactam resistance. **A.** LAC treated with ∏NM1γ6 evolves resistance against ΦNM1γ6, evidenced by the reduction of plaquing from the parental to the evolved populations. **B.** Evolved and parental LAC populations show comparable MICs against different β-lactams and vancomycin.

**Figure S5.**
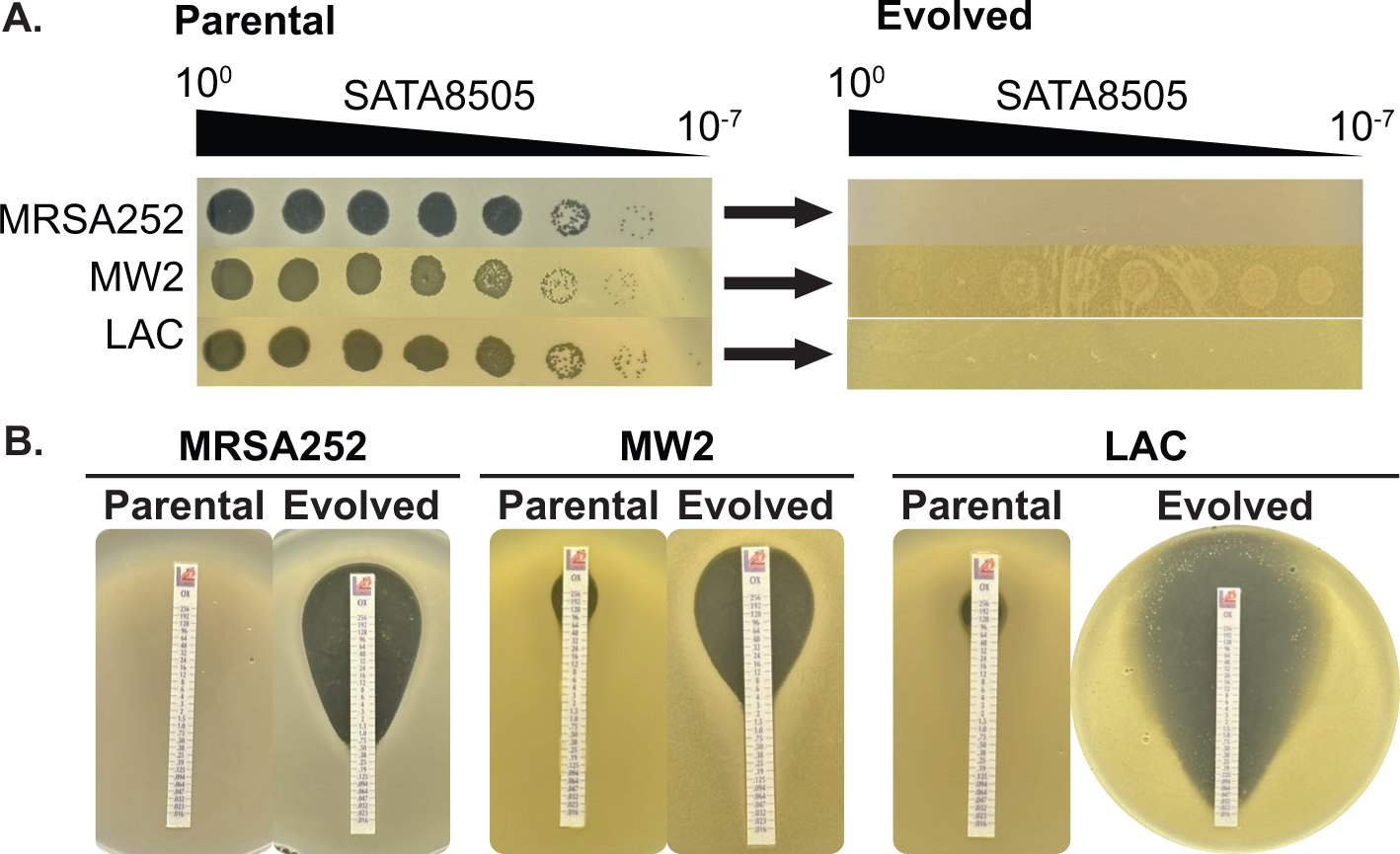
Phage SATA8505 infection drives loss of oxacillin resistance. **A.** MRSA strains MRSA252, MW2, and LAC treated with SATA8585 evolves resistance against SATA8585, evidenced by the reduction of plaquing from the parental to the evolved populations. **B.** Evolved and parental MRSA show reduced MICs against oxacillin.

**Figure S6.**
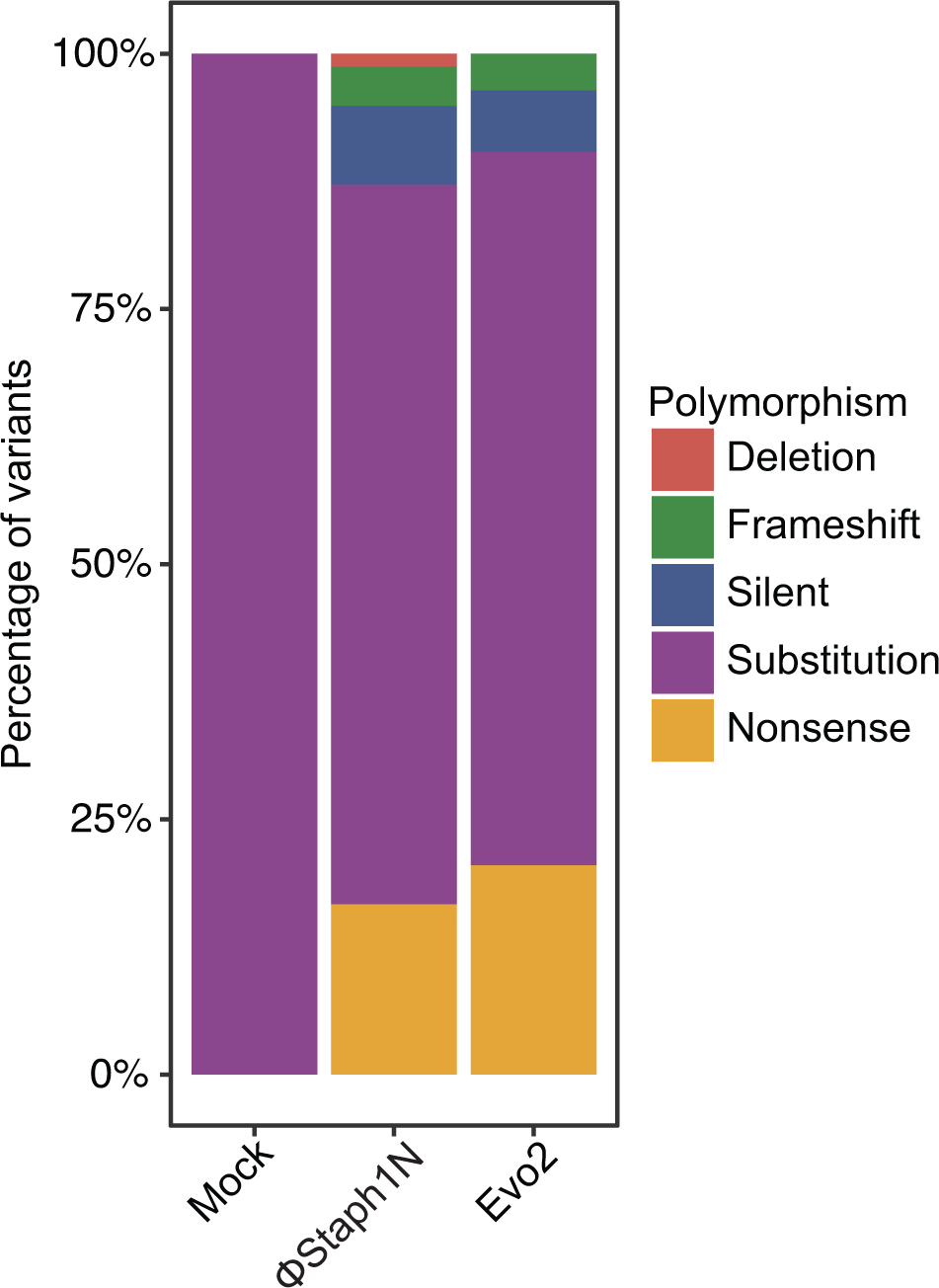
Types of polymorphisms in MRSA strains following infection by phage or a mock treatment. Plotted are the polymorphisms that were found in a gene with an assigned COG category.

**Figure S7.**
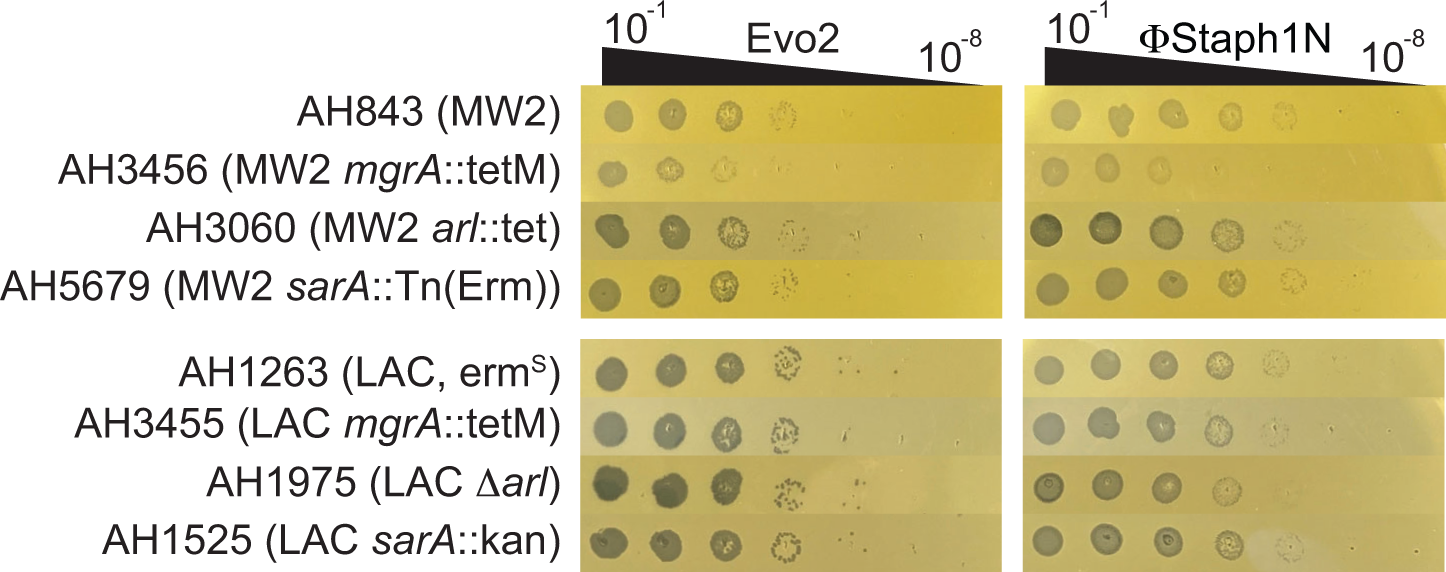
Plaquing efficiency of Evo2 and ΦStaph1N on MW2 and LAC strains with knockouts in *mgrA*, *arl*, and *sarA*.

**Figure S8.**
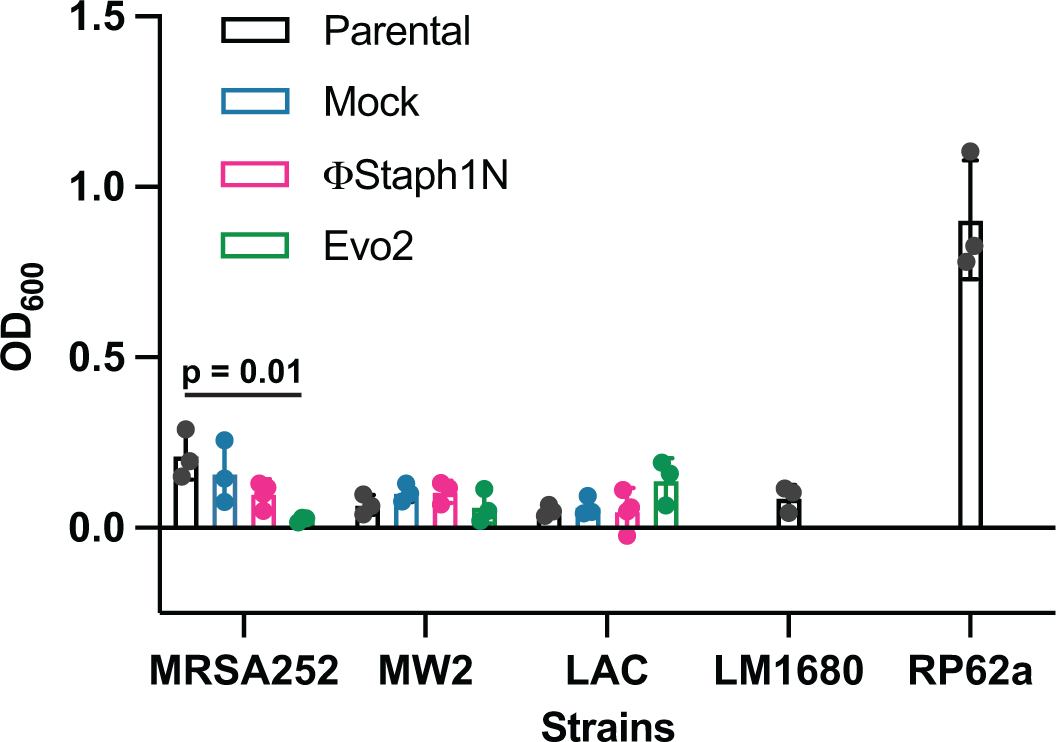
Effect of phage infection on biofilm formation in MRSA strains. Cultures were infected or mock-infected with either ΦStaph1N or Evo2. RP62a is a strain of *S. epidermidis* with known biofilm-forming capability, while LM1680 is a derivative of RP62a that has lost biofilm-forming ability.^69,70^ Biofilm biomass was assessed by staining with Crystal Violet. Solubilized crystal violet was quantified by measuring absorbance at 600 nm. Values represent averages and standard deviations of three replicates. Statistical significance was determined with a two-tailed t-test.

**Table S1.** Bacterial strains and bacteriophages used in this study.

| **Strain Name** | **Comments** | **Source or Reference** |
| --- | --- | --- |
| MRSA MW2 (USA400) |  | Baba *et al.*, 2002 |
| MRSA LAC (USA300) |  | Voyich *et al.*, 2005 |
| MRSA252 (USA200) |  | Holden *et al.*, 2004 |
| *S. epidermidis* RP62a | methicillin-resistant biofilm-producing *S. epidermidis* | Christensen *et al.*, 1982 |
| *S. epidermidis* LAM1680 | Derived from *S. epidermidis* RP62a; carries genomic deletion that inactivates biofilm production | Jiang *et al.*, 2016 |
| ΦStaph1N |  | Łobocka *et al.*, 2012 |
| Evo2 | Derived from ΦStaph1N | This study |
| ΦNM1γ6 | Lytic version of ΦNM1 | Marraffini laboratory |
| ΦNM4γ4 | Lytic version of ΦNM4 | Marraffini laboratory |
| Φ12 |  | Marraffini laboratory |
| Andhra | Infects *S. epidermidis* | Hatoum-Aslan laboratory |
| SATA8505 | Isolated from the environment in this study | Environmental isolate; Pincus *et al.*, 2015 |
| ADL1 | USA300, Patient isolate | Levin laboratory; Land *et al.*, 2015 |
| ADL2 | USA300, Patient isolate | Levin laboratory; Land *et al.*, 2015 |
| ADL3 | USA300, Patient isolate | Levin laboratory; Land *et al.*, 2015 |
| ADL4 | USA300, Patient isolate | Levin laboratory; Land *et al.*, 2015 |
| ADL5 | USA300, Patient isolate | Levin laboratory; Land *et al.*, 2015 |
| ADL6 | USA300, Patient isolate | Levin laboratory; Land *et al.*, 2015 |
| ADL7 | USA300, Patient isolate | Levin laboratory; Land *et al.*, 2015 |
| ADL8 | USA300, Patient isolate | Levin laboratory;<br>Land *et al.*, 2015 |
| ADL9 | USA300, Patient isolate | Levin laboratory;<br>Land *et al.*, 2015 |
| ADL10 | USA300, Patient isolate | Levin laboratory;<br>Land *et al.*, 2015 |
| ADL11 | USA300, Patient isolate | Levin laboratory;<br>Land *et al.*, 2015 |
| ADL12 | USA300, Patient isolate | Levin laboratory;<br>Land *et al.*, 2015 |
| ADL13 | USA300, Patient isolate | Levin laboratory;<br>Land *et al.*, 2015 |
| ADL14 | USA300, Patient isolate | Levin laboratory;<br>Land *et al.*, 2015 |
| ADL15 | USA300, Patient isolate | Levin laboratory;<br>Land *et al.*, 2015 |
| ADL16 | USA300, Patient isolate | Land *et al.*, 2015 |
| ADL17 | USA300, Patient isolate | Levin laboratory;<br>Land *et al.*, 2015 |
| ADL18 | USA300, Patient isolate | Levin laboratory;<br>Land *et al.*, 2015 |
| ADL19 | USA300, Patient isolate | Levin laboratory;<br>Land *et al.*, 2015 |
| ADL20 | USA300, Patient isolate | Levin laboratory;<br>Land *et al.*, 2015 |
| ADL21 | USA300, Patient isolate | Levin laboratory;<br>Land *et al.*, 2015 |
| ADL22 | USA300, Patient isolate | Levin laboratory;<br>Land *et al.*, 2015 |
| ADL23 | USA300, Patient isolate | Levin laboratory;<br>Land *et al.*, 2015 |
| ADL24 | USA300, Patient isolate | Levin laboratory;<br>Land *et al.*, 2015 |
| ADL25 | USA300, Patient isolate | Levin laboratory;<br>Land *et al.*, 2015 |
| ADL26 | USA300, Patient isolate | Levin laboratory;<br>Land *et al.*, 2015 |
| ADL27 | USA300, Patient isolate | Levin laboratory;<br>Land *et al.*, 2015 |
| ADL28 | USA300, Patient isolate | Levin laboratory;<br>Land *et al.*, 2015 |
| ADL29 | USA300, Patient isolate | Levin laboratory;<br>Land *et al.*, 2015 |
| ADL30 | USA300, Patient isolate | Levin laboratory;<br>Land *et al.*, 2015 |
| AH1263 | LAC, Erm<sup>S</sup> | Horswill 850<br>laboratory |
| AH3455 | LAC *mgrA*::tetM | Horswill<br>laboratory |
| AH1975 | LAC Δ*arl* | Horswill<br>laboratory |
| AH1525 | LAC *sarA*::kan | Horswill<br>laboratory |
| AH843 | MW2 | Horswill<br>laboratory |
| AH3456 | MW2 *mgrA*::tetM | Horswill<br>laboratory |
| AH3060 | MW2 *arl*::tet | Horswill<br>laboratory |
| AH5679 | MW2 *sarA*::Tn(Erm) | Horswill<br>laboratory |

**Table S2.** MICs (µg/mL) against oxacillin of MRSA strains treated with different MOIs of phage.

|  | **MRSA252** |  |  |  |  |  |
| --- | --- | --- | --- | --- | --- | --- |
|  | **$\Phi\text{Staph1N}$** |  |  | **Evo2** |  |  |
| **MOI** | **Rep 1** | **Rep 2** | **Rep 3** | **Rep 1** | **Rep 2** | **Rep 3** |
| $10^{-2}$ | 0.25 | 0.125 | 0.38 | NG | 2 | 0.5 |
| $10^{-3}$ | NG | 0.94 | 0.19 | 1 | 0.75 | 1 |
| $10^{-4}$ | 0.5 | 0.25 | 0.19 | 0.75 | 1 | 0.5 |
| $10^{-5}$ | 0.25 | 0.38 | NG | 0.38 | NG | NG |
| Mock | >256 | >256 | >256 | >256 | >256 | >256 |
|  | **MW2** |  |  |  |  |  |
|  | **$\Phi\text{Staph1N}$** |  |  | **Evo2** |  |  |
| **MOI** | **Rep 1** | **Rep 2** | **Rep 3** | **Rep 1** | **Rep 2** | **Rep 3** |
| $10^{-2}$ | 3 | 24 | 24 | 4 | NG | NG |
| $10^{-3}$ | 32 | 24 | 48 | 4 | NG | NG |
| $10^{-4}$ | 48 | 96 | 32 | 3 | NG | NG |
| $10^{-5}$ | 96 | 64 | 24 | 2 | NG | NG |
| Mock | 96 | 48 | 32 | 96 | 48 | 32 |
|  | **LAC** |  |  |  |  |  |
|  | **$\Phi\text{Staph1N}$** |  |  | **Evo2** |  |  |
| **MOI** | **Rep 1** | **Rep 2** | **Rep 3** | **Rep 1** | **Rep 2** | **Rep 3** |
| $10^{-2}$ | NG | NG | 2 | 0.064 | NG | NG |
| $10^{-3}$ | NG | 3 | 1.5 | 0.032 | NG | NG |
| $10^{-4}$ | 32 | 1.5 | 1 | NG | NG | NG |
| $10^{-5}$ | 32 | 16 | 0.38 | NG | NG | NG |
| Mock | 32 | 48 | 48 | 32 | 48 | 48 |

NG: no growth detected

**Table S3.** MICs (µg/mL) of mock- or Evo2-treated MRSA strains against different antibiotics.

| Strain | Antibiotic | Mock |  |  | Evo2 |  |  |
| --- | --- | --- | --- | --- | --- | --- | --- |
|  |  | Rep 1 | Rep 2 | Rep 3 | Rep 1 | Rep 2 | Rep 3 |
| MRSA252 | Oxacillin | >256 | >256 | >256 | 0.38 | 0.75 | 0.5 |
|  | Rifampicin | 0.047 | 0.032 | 0.012 | 0.023 | 0.047 | 0.023 |
|  | Mupirocin | 0.75 | 0.5 | 1 | 0.5 | 0.5 | 0.38 |
|  | Erythromycin | >256 | >256 | >256 | >256 | >256 | >256 |
|  | Teicoplanin | 6 | 6 | 4 | 4 | 4 | 0.75 |
|  | Fosfomycin | 12 | 8 | 8 | 8 | 8 | 6 |
|  | Daptomycin | 2 | 3 | 2 | 2 | 2 | 2 |
| MW2 | Oxacillin | 48 | 32 | 32 | 0.75 | 1 | 0.75 |
|  | Rifampicin | 0.032 | 0.047 | 0.064 | 0.023 | 0.023 | 0.032 |
|  | Mupirocin | 0.5 | 0.25 | 0.5 | 0.38 | 0.38 | 0.25 |
|  | Erythromycin | 1 | 0.75 | 0.75 | 0.25 | 0.5 | 0.25 |
|  | Teicoplanin | 2 | 1.5 | 1.5 | 1.5 | 1 | 1 |
|  | Fosfomycin | 1 | 1.5 | 1.5 | 0.5 | 1 | 1 |
|  | Daptomycin | 1.5 | 2 | 1.5 | 0.75 | 0.5 | 3 |
| LAC | Oxacillin | 48 | 24 | 64 | 0.19 | 0.064 | 0.047 |
|  | Rifampicin | 0.047 | 0.047 | 0.047 | 0.032 | 0.032 | 0.047 |
|  | Mupirocin | 0.5 | 0.75 | 0.75 | 0.5 | 0.5 | 0.5 |
|  | Erythromycin | 3 | 3 | 3 | 1.5 | 1 | 2 |
|  | Teicoplanin | 1 | 0.5 | 1 | 0.5 | 0.5 | 0.75 |
|  | Fosfomycin | 6 | 6 | 12 | 1.5 | 12 | 1 |
|  | Daptomycin | 0.5 | 3 | 1 | 0.064 | 2 | 0.75 |

**Table S4.**
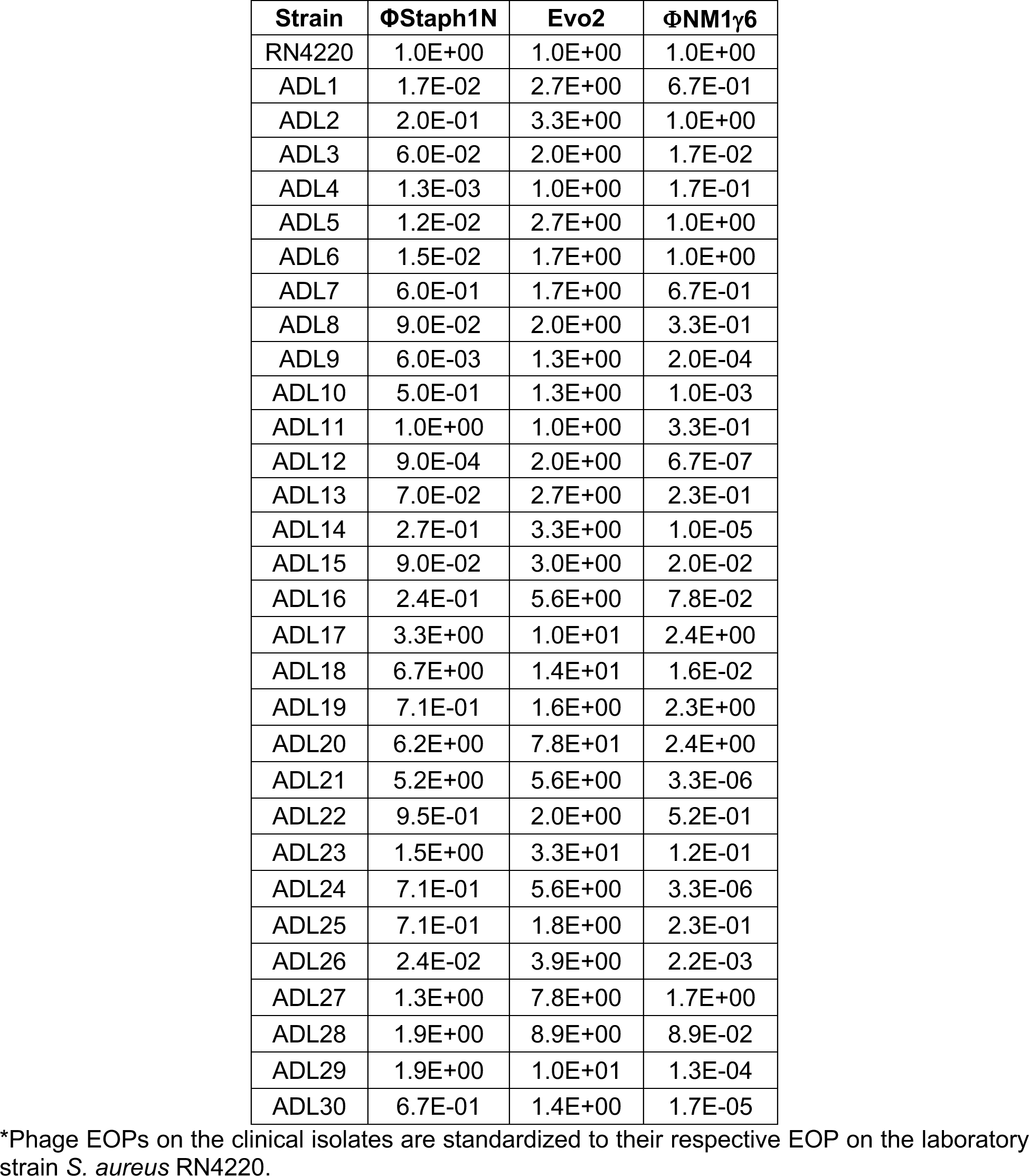
Efficiencies of plaquing (EOPs)* of ΦStaph1N, Evo2, and ∏NM1γ6 on clinical isolates of USA300 (ADL1-30).

**Table S5.**
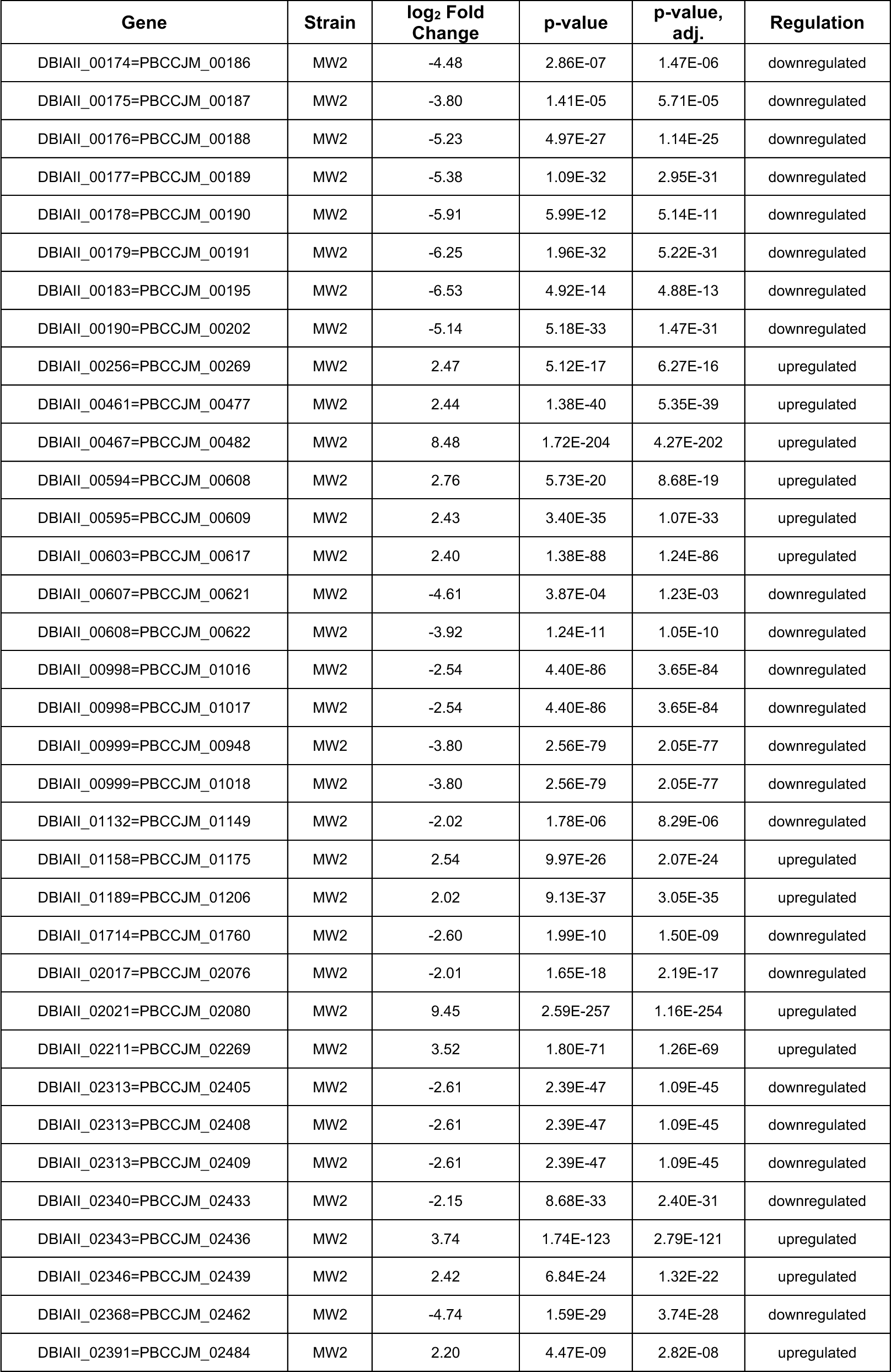
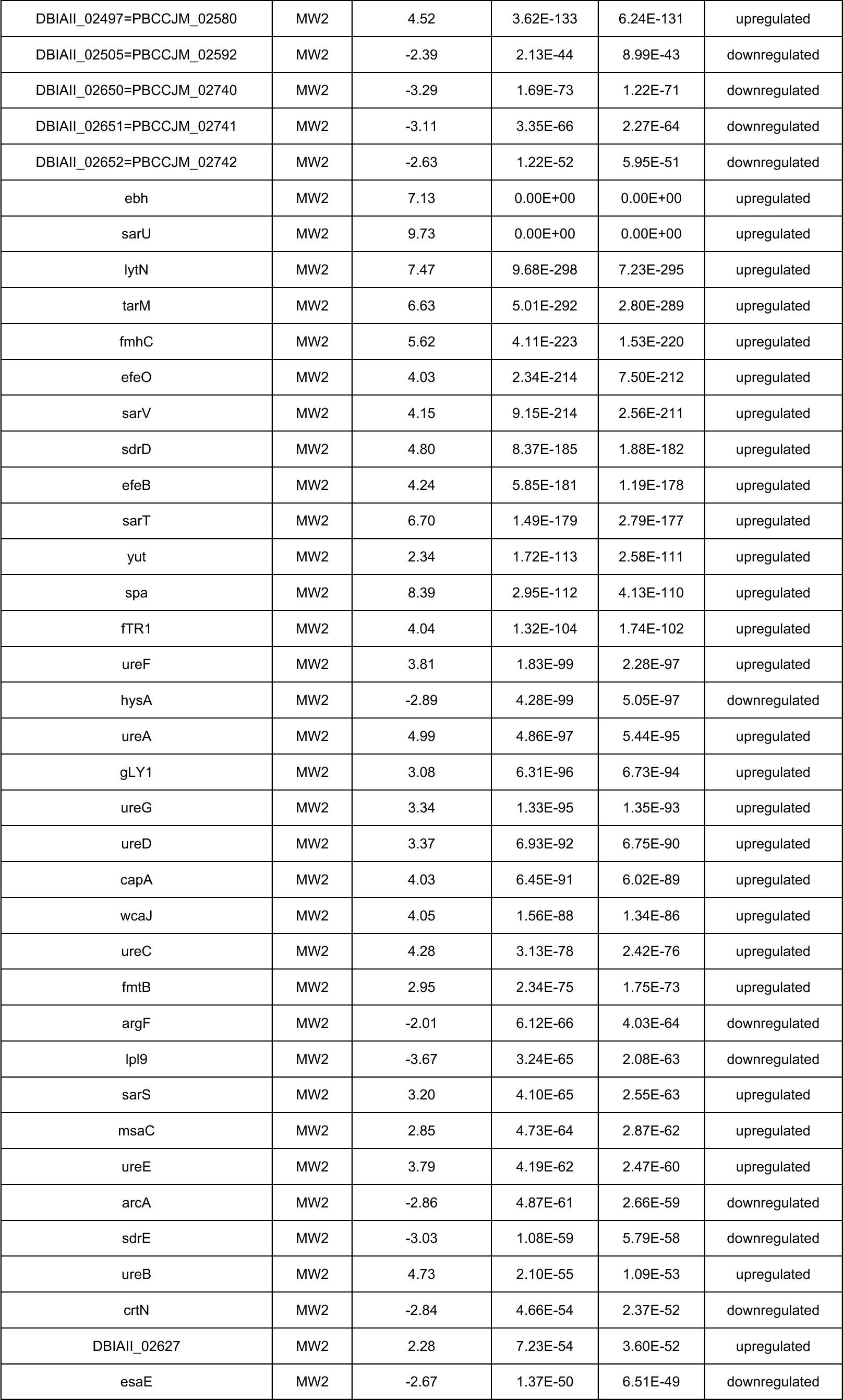
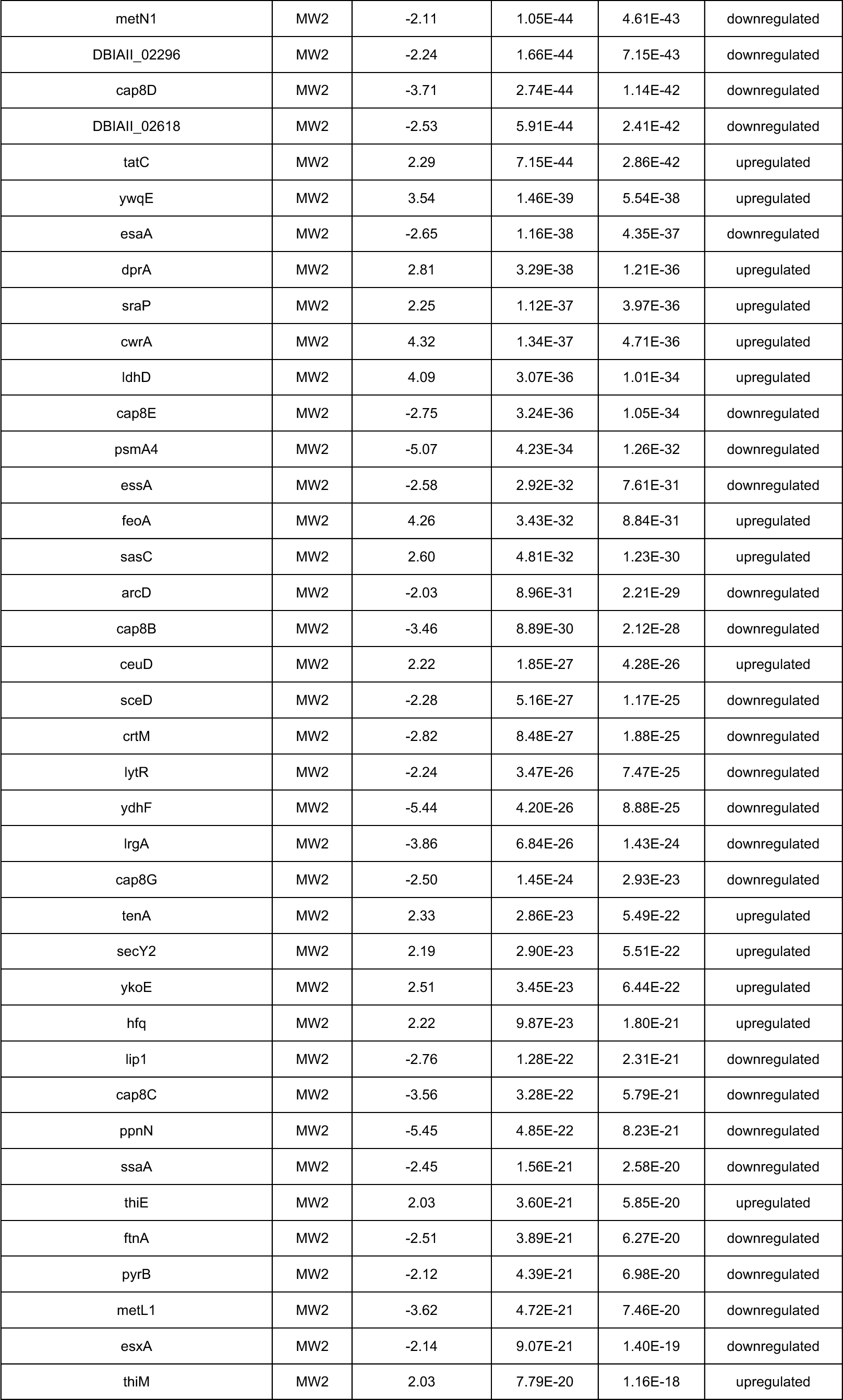
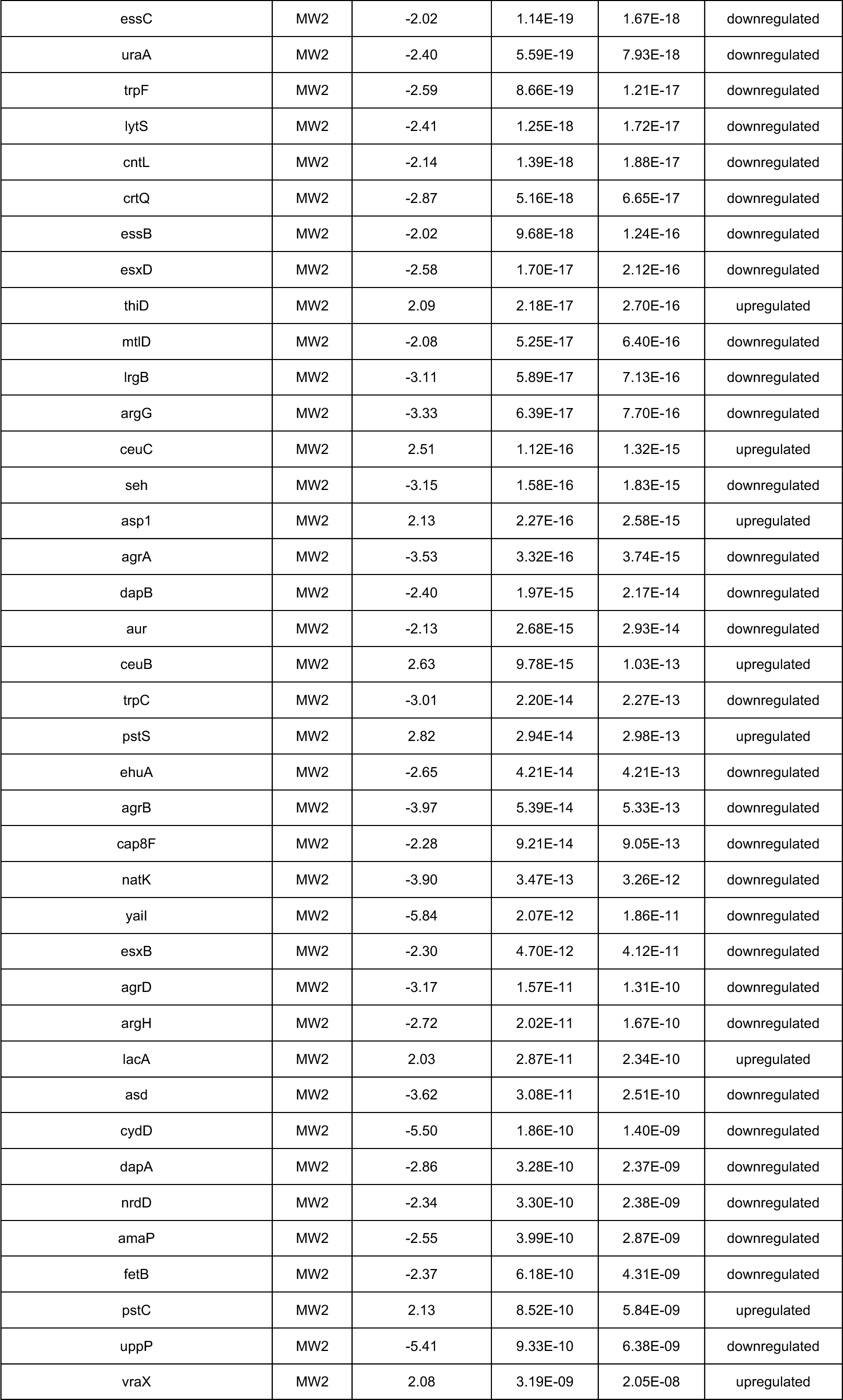
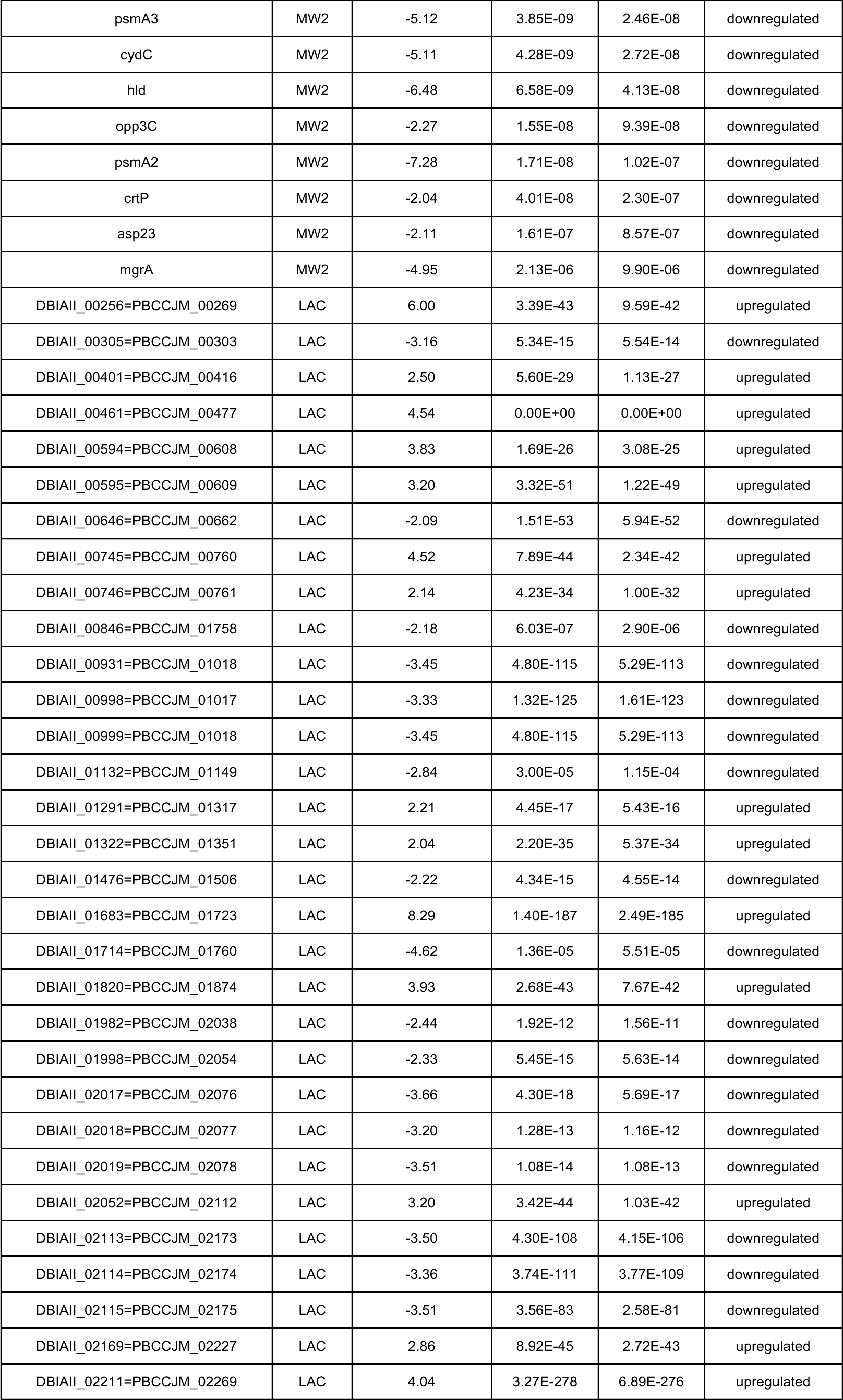
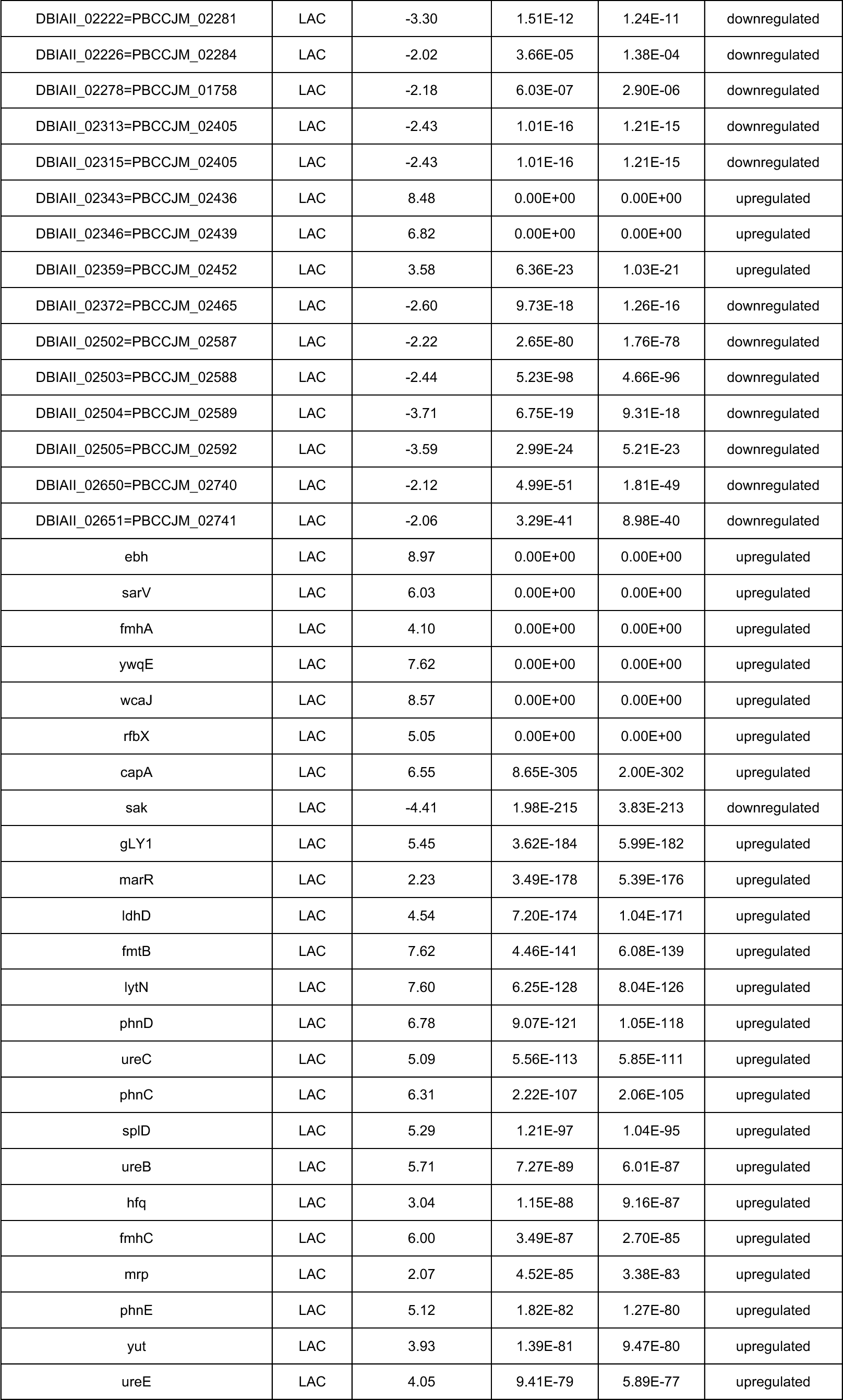
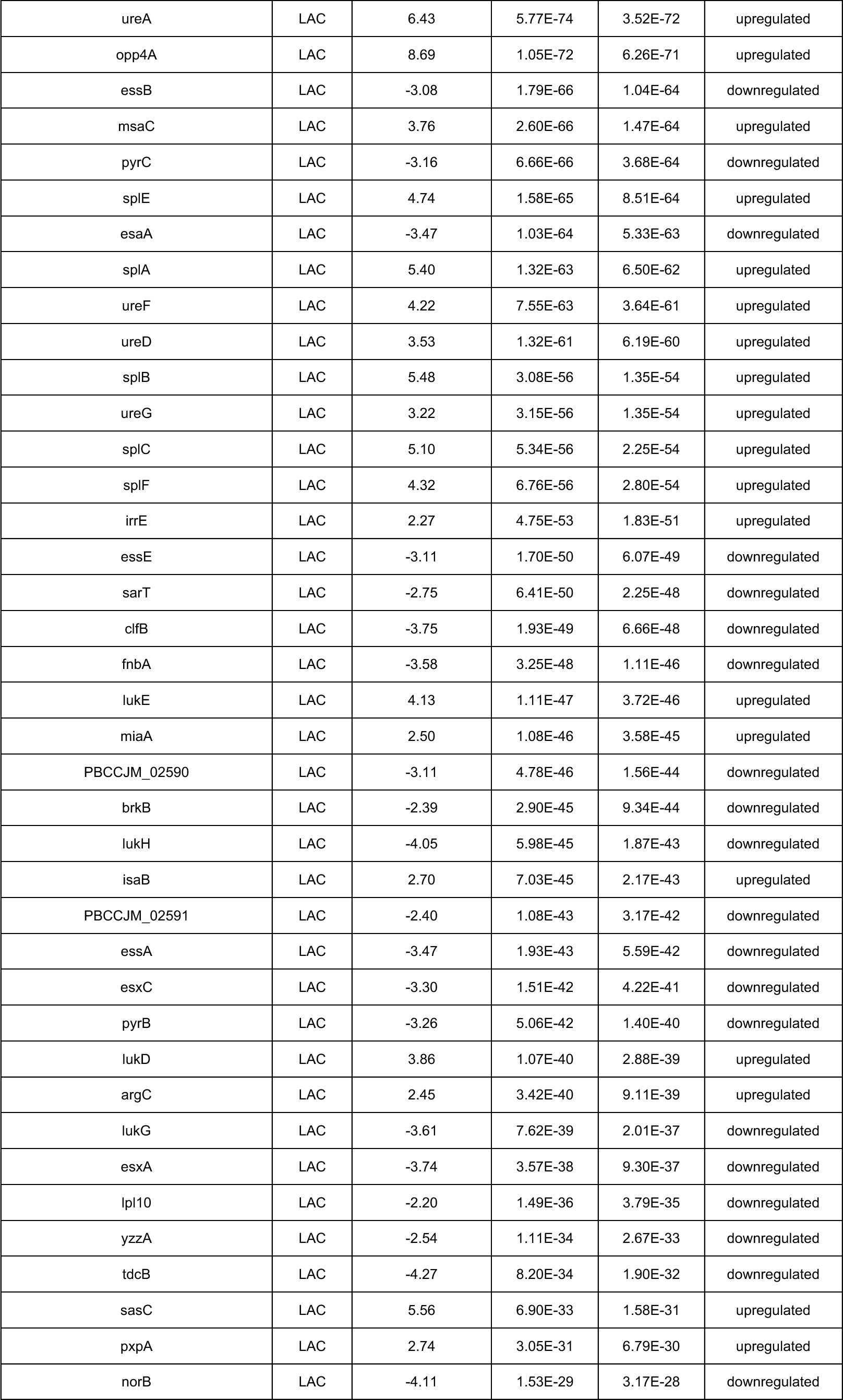
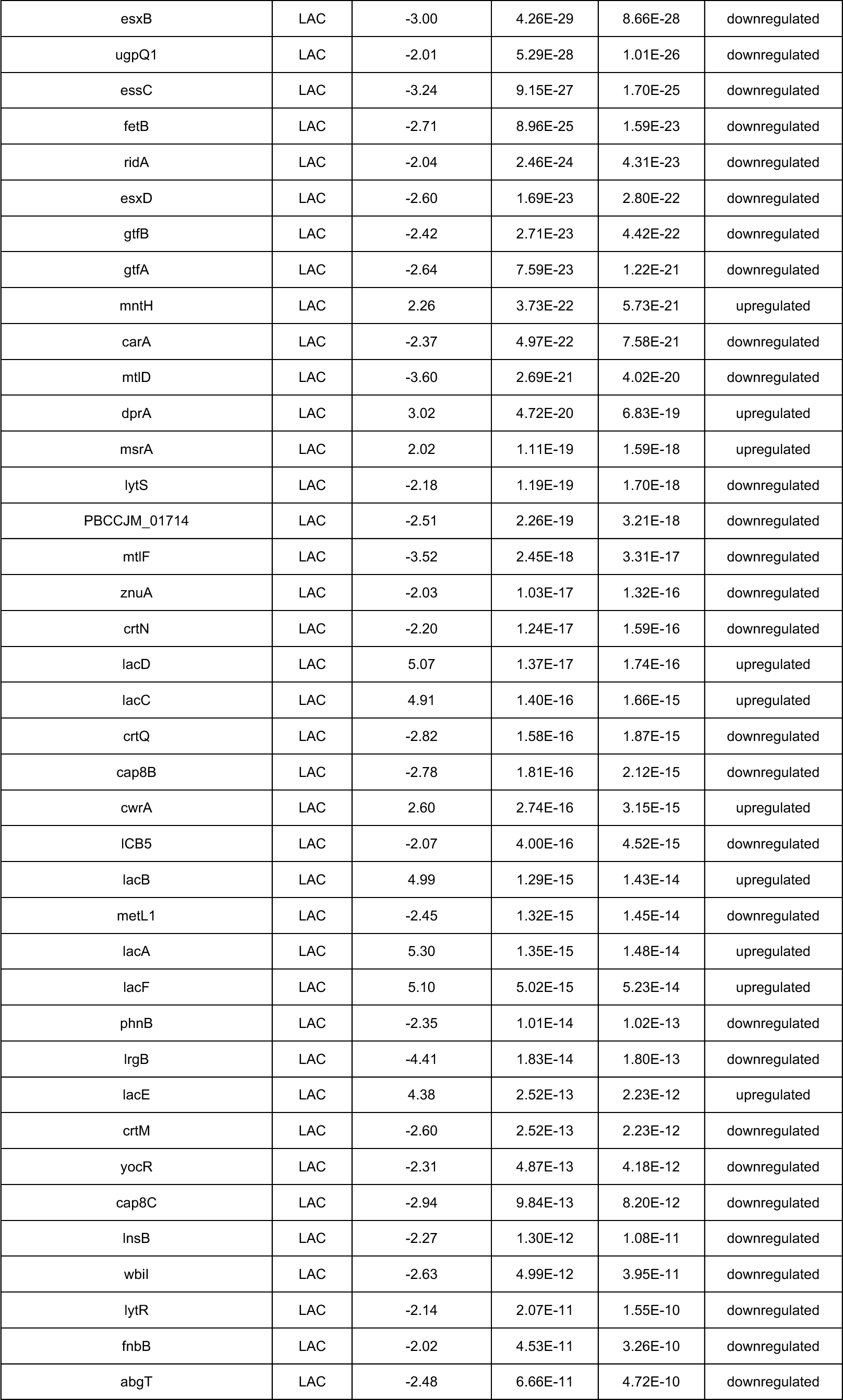
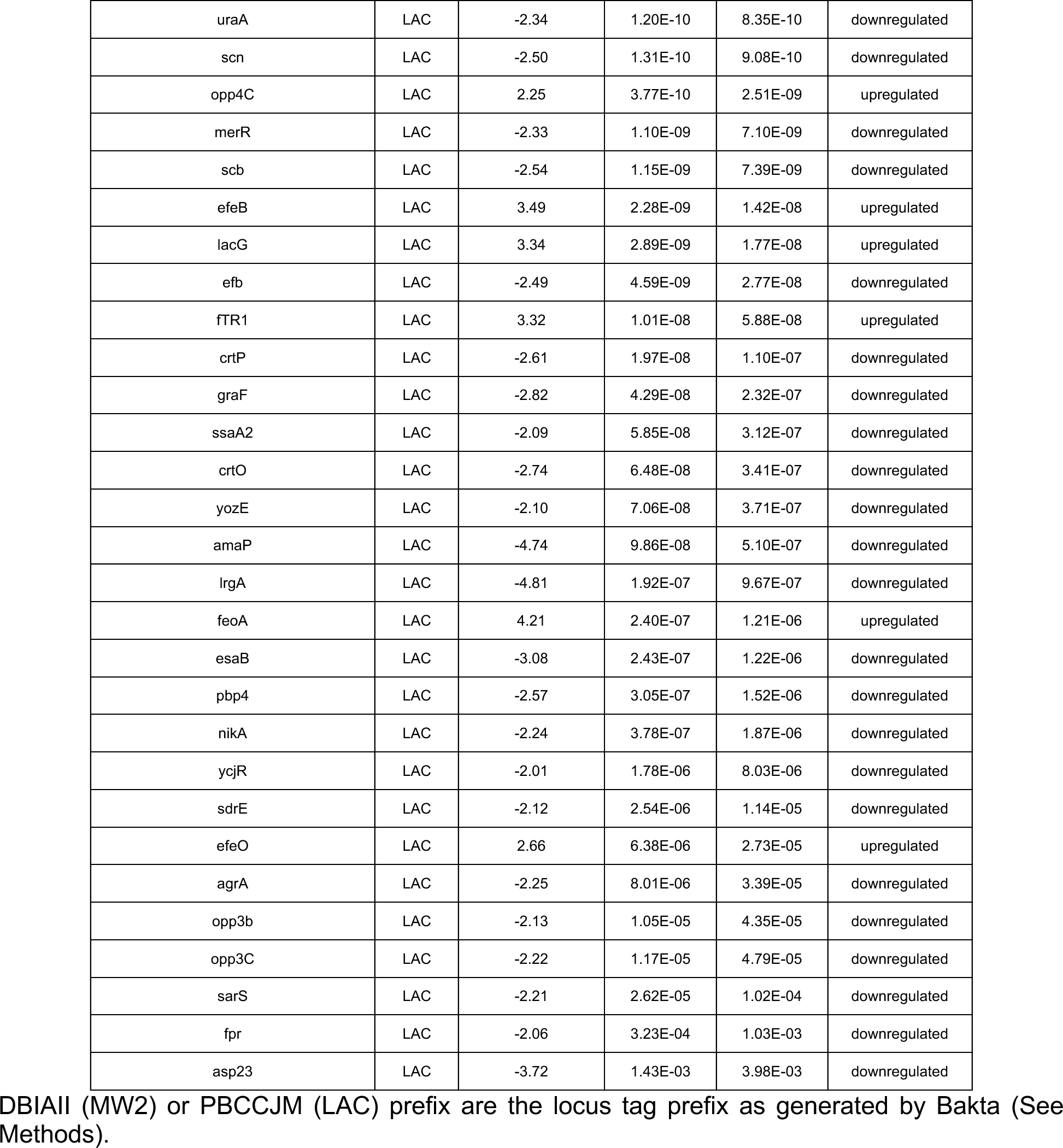
Genes with significant fold changes in MRSA MW2 and LAC strains following Evo2 infection.

**Table S6.** Mutations in surviving MW2 cells co-treated with ΦStaph1N and oxacillin.

| Gene | Description | Reference |
| --- | --- | --- |
| rsaC<br>ncRNA | modulate oxidative stress response and metal immunity | https://doi.org/10.1093/nar/gkz728 |
| nrdF | class 1b ribonucleoside-diphosphate reductase subunit beta; beta subunit contains a metal-based cofactor; involved in DNA synthesis | [10.1128/JB.183.24.7260-7272.2001](https://doi.org/10.1128/JB.183.24.7260-7272.2001)<br>10.1074/jbc.M113.533554 |
| fstAT<br>ncRNA | Unknown |  |
| rpoC | DNA-directed RNA polymerase subunit beta' |  |
| tRNA | Transfer RNA |  |

